# eIF4E3 Forms an Active eIF4F Complex during Stresses (eIF4F^S^) Targeting mTOR and Re-Programs the Translatome

**DOI:** 10.1101/2021.01.22.427788

**Authors:** B. Weiss, G.E. Allen, J. Kloehn, K. Abid, P. Jaquier-Gubler, J.A. Curran

**Affiliations:** Department of Microbiology and Molecular Medicine, Faculty of Medicine, University of Geneva, Switzerland; Catecholamine and Peptides Laboratory, Service of Clinical Pharmacology, Lausanne University Hospital and University of Lausanne, Switzerland; Institute of Genetics and Genomics of Geneva (iGE3), University of Geneva, Switzerland

## Abstract

The eIF4E are a family of initiation factors that bind the mRNA 5’ cap, regulating the proteome and the cellular phenotype. eIF4E1 mediates global translation and its activity is controlled via the PI3K/AKT/mTOR pathway. mTOR down-regulation results in eIF4E1 sequestration into an inactive complex with the 4E binding proteins (4EBPs). The second member, eIF4E2, regulates the translatome during hypoxia. However, the exact function of the third member, eIF4E3, has remained elusive. We have dissected its function using a range of techniques. Starting from the observation that it does not interact with 4EBP1, we demonstrate that eIF4E3 recruitment into an eIF4F complex occurs when Torin1 inhibits the mTOR pathway. Ribo-seq studies demonstrate that this complex (eIF4F^S^) is translationally active only during stress and that it selects specific mRNA populations based on 5’ TL (UTR) length. The interactome reveals that it associates with cellular proteins beyond the cognate initiation factors, suggesting that it may have “moon-lighting” functions. Finally, we provide evidence that cellular metabolism is altered in an eIF4E3 KO background but only upon Torin1 treatment. We propose that eIF4E3 acts as a second branch of the integrated stress response, re-programming the translatome to promote “stress resistance” and adaptation.

## Introduction

Translation is the most energy consuming process in the cell and is subjected to regulation (Pannevis and Houlihan, 1992). Most of this is exerted at the initiation step impacting on the proteome, and as a consequence, the cellular phenotype (Hinnebusch et al., 2016). Initiation involves the loading of the 40S ribosomal subunit onto the mRNA 5’ end and subsequent scanning of the 5’ transcript leader (5’ TL or 5’ UTR) (Curran and Weiss, 2016; Hinnebusch and Lorsch, 2012). This process engages a plethora of eukaryotic translation initiation factors (eIFs) that associate both with the 40S and the mRNA 5’ end. The 40S interacts with the scaffolding protein eIF3, the ternary complex eIF2.GTP.tRNA^MET^, eIF1/1A and the eIF5 GTPase. Together they form the 43S preinitiation complex (PIC) that loads onto the mRNA via an eIF3-eIF4G interaction. eIF4G is a member of the eIF4F trimolecular complex that assembles on the 5’ cap and carries the eIF4E cap-binding protein and the eIF4A DEAD-box helicase (Merrick, 2015). Once loaded, the PIC scans the 5’ TL until an AUG codon is recognized. This triggers eIF2.GTP hydrolysis, release of the 40S associated initiation factors and the recruitment of the 60S subunit to generate the functional 80S ribosome. The eIF2.GDP generated at each round of initiation is recycled back into its active GTP form by the eIF2B exchange factor (Hinnebusch, 2012).

The cellular response to a multitude of stresses impacts the translatome by targeting either eIF4E or eIF2. The formation of the eIF4F complex can be disrupted by the 4E binding proteins (4EBPs) that compete with eIF4G for eIF4E because of their common YxxxxLΦ (Φ being hydrophobic) interacting motif. The 4EBPs are phosphoproteins and their affinity for eIF4E responds to phosphorylation status (Gingras et al., 1998), with hypophosphorylated 4EBPs having strong affinity and hyperphosphorylated forms being unable to interact (Gingras et al., 2001). Phosphorylation is regulated by the mechanistic target of rapamycin complex 1 (mTORC1), a kinase conserved from yeast to human (Loewith et al., 2002). The core mTOR kinase actually exists in two functionally distinct complexes, mTORC1 and mTORC2 (Kim et al., 2017). The former influences cell growth by targeting downstream effectors that regulate protein translation, including not only the 4EBPs but also the ribosomal protein S6 kinase 1 (S6K1) (Sabatini, 2006; Volarević and Thomas, 2001). It is responsive to the PI3K/AKT pathway and to intracellular ATP, glucose, and amino acid levels (Loewith et al., 2002). The mTORC2 kinase activity is coupled to numerous extracellular signalling pathways and has been linked to cytoskeletal organization and cell survival (Jacinto et al., 2004). Furthermore, the mTORC1 and mTORC2 auto-regulate each other (Jhanwar-Uniyal et al., 2019). The eIF2 activity is regulated by a family of stress activated protein kinases that phosphorylate the eIF2α subunit of eIF2.GDP (Baird and Wek, 2012). The P-eIF2.GDP acts as a competitive inhibitor of eIF2B and because the eIF2 concentration is higher than eIF2B, phosphorylation of even a fraction of eIF2 can provoke a rapid fall in the active eIF2B cellular levels (Hinnebusch, 2005). Consequently, TC levels fall, affecting negatively global protein synthesis. This pathway is referred to as the integrated stress response (ISR) (Pakos-Zebrucka et al., 2016).

eIF4E is a family of cap binding proteins that consists of three members in mammals (Ho and Lee, 2016; Joshi et al., 2004). eIF4E1 is the prototype of the family and is essential. It has the highest affinity for the 5’ cap, interacting through two tryptophan residues forming an aromatic sandwich around the 5’ m^7^Gppp (Osborne et al., 2013). It is a proto-oncogene found over-expressed in 30% of human tumours (Graff and Zimmer, 2003; Topisirovic et al., 2003). Its oncogenic properties are coupled to its ability to shuttle between cytoplasm and nucleus as a 4E-mRNA transporter (Culjkovic-Kraljacic et al., 2012; Dostie et al., 2000). The other two members of the family are eIF4E2 and eIF4E3. eIF4E2 interacts with the 4EBP proteins and is essential for development as knock-out (KO) mice show perinatal lethality (Morita et al., 2012). This was attributed to the selective translational inhibition of mRNA subpopulations (Morita et al., 2012). In the adult, it does not appear to play a major role in translation, as it cannot interact with eIF4G1 (Cho et al., 2006; Joshi et al., 2004; Morita et al., 2012). However, in hypoxia, it is recruited onto mRNAs through the hypoxia inducible factor HIF1α forming an eIF4F complex with eIF4G3 (Uniacke et al., 2012; Uniacke et al., 2014). This eIF4F^H^ complex re-programs the translational readout in response to hypoxia-induced stress. The last member of the family, eIF4E3 is non-essential, as the KO mice are viable (IMPC: https://www.mousephenotype.org). The crystal structure of eIF4E3-m7Gppp revealed that it interacts with the cap through a single tryptophan and a hydrophobic side chain (Osborne et al., 2013). The interaction is 40 fold weaker than that formed by eIF4E1. Over-expression studies indicated that eIF4E3 interacted with eIF4G1 and eIF4G3 but not 4EBP (Frydryskova et al., 2016; Joshi et al., 2004; Landon et al., 2014). Unlike eIF4E1, eIF4E3 over-expression does not transform cells, however, it does down-regulate the translation of transcripts up regulated upon eIF4E1 over-expression. This led the authors to propose that it is a tissue-specific tumour suppressor (Osborne et al., 2013). An expression profile comparison on cells over-expressing either eIF4E1 or eIF4E3 revealed only modest changes in the translatome with the most significant changes mapping to the transcriptome (Landon et al., 2014). Reports have also implicated eIF4E3 in the progression of prostate and medulloblastoma cancers (Abdelfattah et al., 2018; Lin et al., 2007).

In this manuscript, we have dissected the function of eIF4E3. We demonstrate that it forms a functional eIF4F complex only during Torin1 induced stress. This is translationally active selecting transcripts based upon 5’ TL length. We have probed the interactome of eIF4E3 establishing that it associates with proteins beyond the cognate initiation factors, suggesting “moon-lighting” functions. Finally, we provide evidence that cellular metabolism is compromised in an eIF4E3 KO background but only under Torin1-induced stress conditions.

## RESULTS

### eIF4E3 binds the cap and is not regulated by 4EBP1

eIF4E3 has cap binding activity (Osborne et al., 2013). However, the interaction involves only a single tryptophan residue (W115 in the human protein), whereas eIF4E1 traps the cap between two tryptophans creating an aromatic sandwich (W56 and W110 in the murine protein). To confirm the role of W115, HA-tagged versions of wild type and mutant murine eIF4E1 and human eIF4E3 were transiently overexpressed. Cap binding proteins were selected and characterised by Western blotting (Figure 1A and 1B). While we could pull-down WT forms of eIF4E1^HA^ and eIF4E3^HA^, none of the eIF4E1^HA^ single tryptophan mutants (W56A and W110A) or eIF4E3^HA^ W115A were retained.

**Figure 1.**
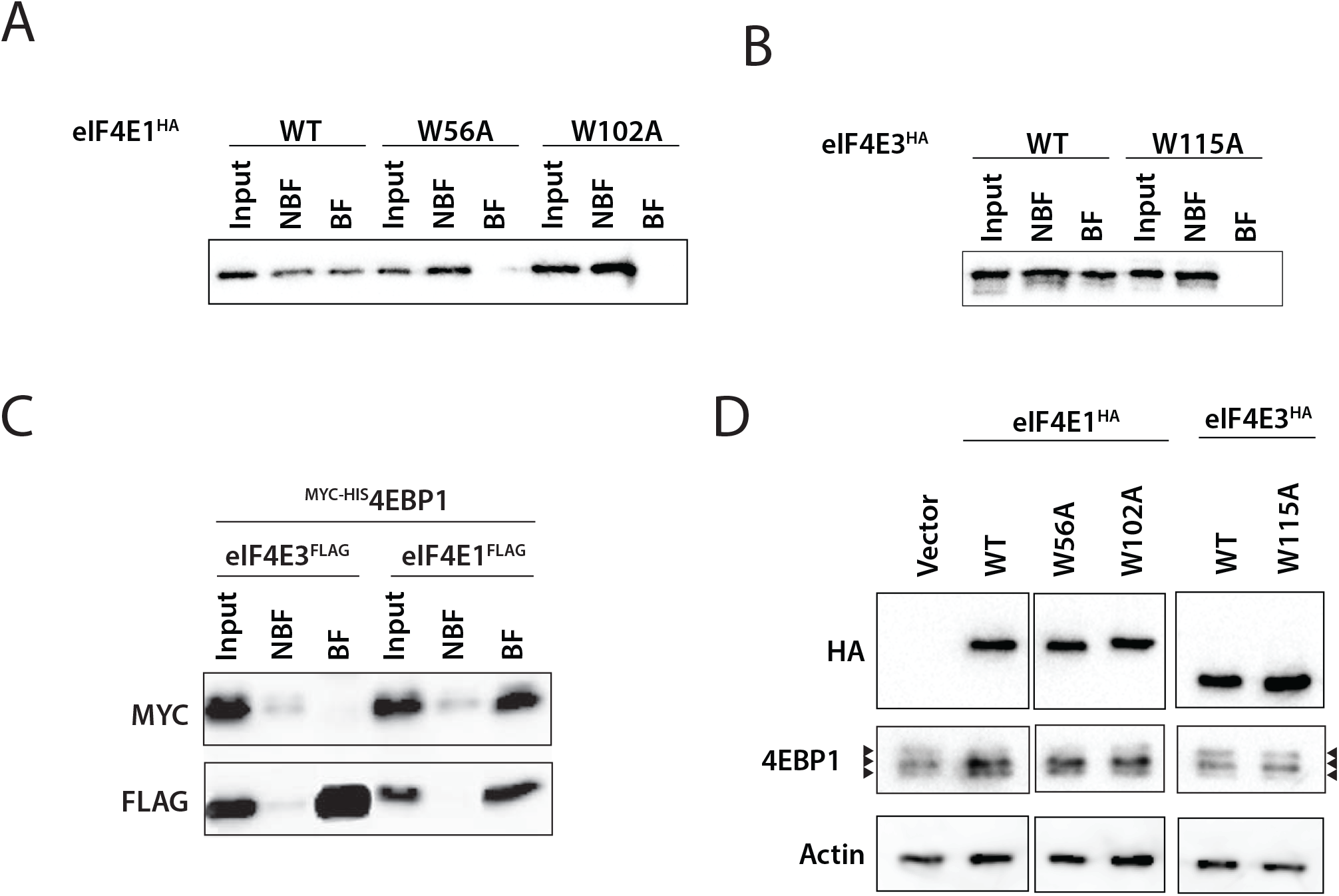
eIF4E3 binds the 5’ cap and is not regulated by 4EBP1. **A-B**. Anti-HA Western blots following cap pull down using cell lysates from HEK293T cells transfected with WT or tryptophan mutants of eIF4E1^HA^ (**A**) or eIF4E3^HA^ (**B**). The input, non-binding fraction (NBF) and binding fractions (BF) are indicated. **C**. Co-immunoprecipitation assays using either eIF4E3^FLAG^ or eIF4E3^FLAG^ transiently co-expressed with ^HIS-MYC^4EBP1 in HEK293T cells. The HIS-tagged protein was pulled down and the presence of the FLAG and MYC tagged proteins was monitored by Western blotting. **D**. Western blot showing the phosphorylation status of 4EBP1 in HEK293T cells transduced with empty vector, eIF4E1^HA^ WT and tryptophan mutants or eIF4E3^HA^ WT and tryptophan mutant.

The interaction between eIF4E3 and 4EBP1 was probed by transient co-overexpression of ^MYC/HIS^4EBP1 with eIF4E3^FLAG^ or eIF4E1^FLAG^. eIF4E1^FLAG^ but not eIF4E3^FLAG^ was pulled down with ^MYC/HIS^4EBP1 (Figure 1C), confirming previous observations that had employed purified proteins (Joshi et al., 2004). We noted that whereas 4EBP1 was hypophosphorylated in cells transduced with eIF4E1^HA^ as reported (Yanagiya et al., 2012), it remained largely hyperphosphorylated in cells with empty vector or eIF4E3^HA^ (Figure 1D). The changes in 4EBP1 phosphorylation status were not affected by any of the mutants. These results demonstrate that eIF4E3 is not being regulated by 4EBP1.

### In normal growth conditions, the eIF4F complex carries mainly eIF4E1

Over-expression/co-IP studies indicated that eIF4E3 could form an eIF4F(3) complex (Frydryskova et al., 2016; Joshi et al., 2004; Landon et al., 2014). However, we were unable to reproduce these results with endogenous proteins. Therefore, to follow recruitment of the endogenous eIF4E3 into eIF4F complexes we employed glycerol gradients (Dieudonne et al., 2015; Legrand et al., 2015). Before loading, the cell extracts were divided into two equal fractions. One was heated in denaturing conditions (+SDS) to disrupt complexes. Gradient analysis of native cell extracts revealed that most of the endogenous eIF4E1 co-sedimented in the lower end of the gradient, namely fractions 3/4, with eIF4G1 and eIF4A1 (Figure 2A). A minor fraction co-sedimented with hypophosphorylated 4EBP1 (fraction 7) but not with hyperphosphorylated 4EBP1 (fractions 8-10). eIF4A1, which is a very abundant cellular protein (Beck et al., 2011), sedimented in multiple fractions. With regards to eIF4E3, the vast majority of the protein sedimented in the upper half of the gradient from fractions 5-9, with only a minor amount co-sedimenting with eIF4G1 in fraction 4. Denaturation disrupted all the complexes in the initiation process as confirmed by the shift of both eIF4E1 (the eIF4F complex) and eIF2α (the TC) to the upper fractions (Figure S1) (Legrand et al., 2015). These results suggest that under normal growth conditions eIF4F is composed of eIF4E1, eIF4G1 and eIF4A1, with eIF4E3 remaining in the light pool. We next explored if this eIF4F complex is associated with ribosomes by examining the sedimentation profile of ribosomal protein RPS6. We did not find RPS6 within the gradients from the native extracts (Figure 2B). However, it was detected in the SDS-denatured extracts suggesting that under native conditions it was associated with heavier complexes that had pelleted. This we confirmed by re-suspending the pellets in Laemmli buffer. Western blotting revealed the presence of RPS6, eIF4E1, eIF4G1 and eIF4A1 but not eIF4E3 (Figure 2C). Therefore, our sedimentation analysis monitored the formation of complexes not associated with the 40S.

**Figure 2.**
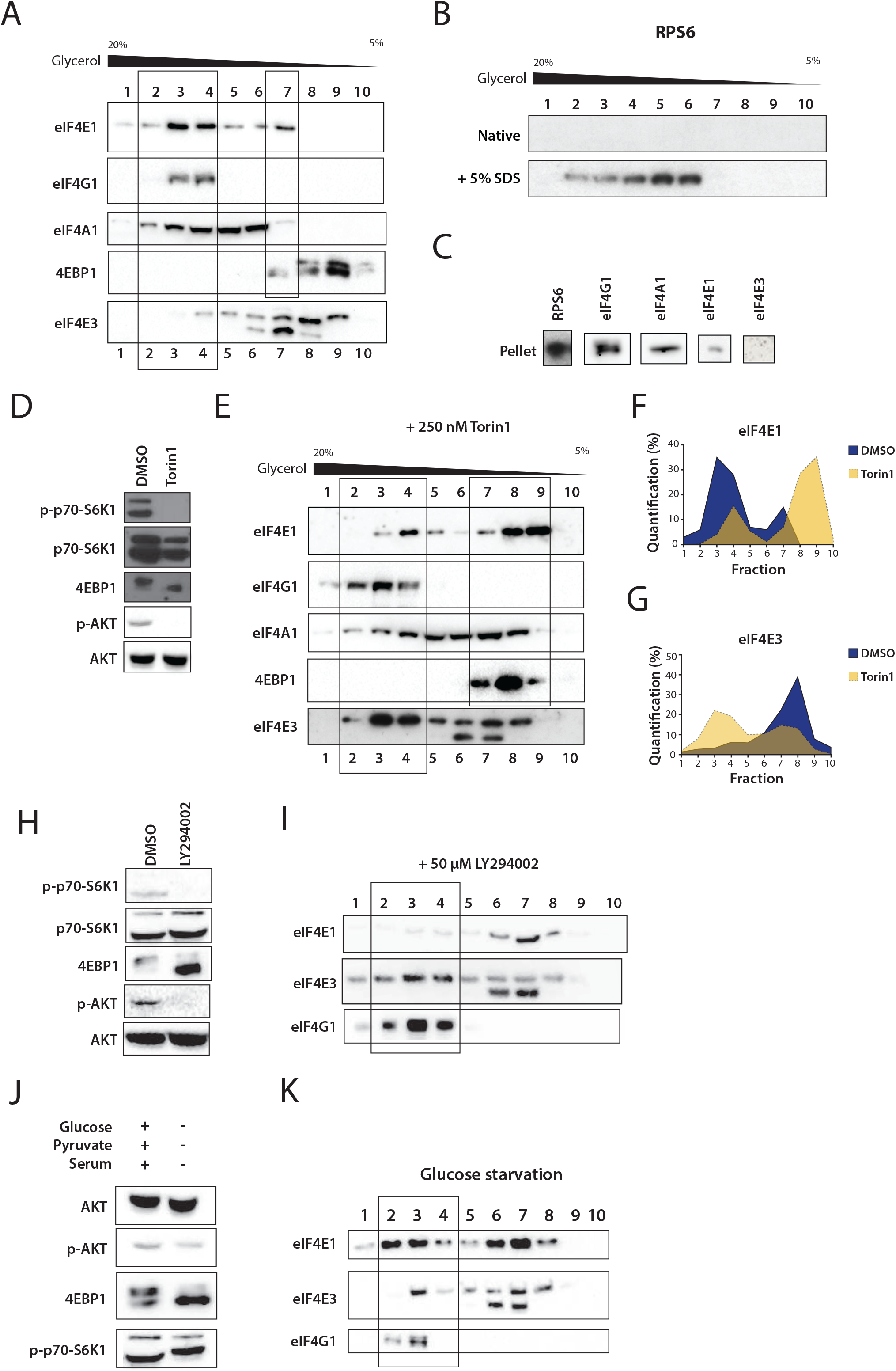
eIF4E3 co-sediments with eIF4F when the AKT/mTOR pathway is inhibited. **A**. Western blots showing the sedimentation profiles on glycerol gradients of members of the eIF4F complex and of 4EBP1 from HEK293T cell lysates. Fractions 2/3/4 and fraction 7 are highlighted as the regions in which we find eIF4F (based upon the sedimentation of eIF4G1) and eIF4E3, respectively. **B**. Western blots of the sedimentation profiles on glycerol gradients of RPS6 from HEK293T cells lysates treated or non-treated with SDS prior to loading. **C**. Western blots of eIF4F complex members following resuspension of the gradient pellet. **D**. Western blots showing the phosphorylation levels of AKT/mTOR pathway members following treatment of HEK293T cells with vehicle or with 250 nM Torin1 for 2 hours. **E**. Western blots showing the sedimentation profiles of members of the eIF4F complex and of 4EBP1 from a lysate of HEK293T cells treated with 250 nM Torin1 for 2 hours. Fractions 2/3/4 and fractions 7/8/9 are highlighted as the regions in which we find eIF4F and the majority of eIF4E1, respectively. **F-G**. Quantification of the level of eIF4E1 (**F**) and eIF4E3 (**G**) derived from the profiles in panels (**A)** and **(E)**. **H**. Western blots showing the phosphorylation status of AKT/mTOR pathway members following treatment of HEK293T cells with vehicle or with 50 μM LY294002 for 1 hour. **I**. Western blots showing the sedimentation profiles of members of the eIF4F complex and 4EBP1 from a lysate of HEK293T cells treated with 50 μM LY294002 for 1 hour. The fractions containing the eIF4F complex are indicated by the rectangle. **J**. Western blots showing the phosphorylation status of AKT/mTOR pathway members following glucose starvation of HEK293T cells for 2 hours. **K**. Western blots showing the sedimentation profiles of members of the eIF4F complex and 4EBP1 from a lysate of HEK293T cells glucose starved for 2 hours. The fractions in which the eIF4F complex is located are indicated by the rectangle.

### eIF4E3 sediments with the eIF4F complex when eIF4E1 is sequestered by 4EBP1

Our results confirm that eIF4E3 does not bind 4EBP1. It also does not co-sediment with eIF4G1 in normal growth conditions. We hypothesised that eIF4E3 would assemble an eIF4F complex under conditions in which eIF4E1 was sequestered by 4EBP1. We therefore treated cells with the mTOR inhibitor Torin1 (Thoreen et al., 2009). This drug targets both mTORC1 and mTORC2 as confirmed by the dephosphorylation of 4EBP1, RPS6K and AKT-Ser^473^ (Figure 2D). We repeated the sedimentation analysis on extracts prepared from cells treated with Torin1 or with vehicle for 2 hours. As expected, a major fraction of eIF4E1 moved from the bottom to the top of the gradient and co-sedimented with hypophosphorylated 4EBP1 (Figure 2E, fractions 7-9). eIF4G1 remained in fractions 2-4. A significant fraction of eIF4E3 now co-sedimented with eIF4G1 in fractions 3 and 4. Therefore, upon mTOR inhibition the sedimentation profiles of eIF4E1 and eIF4E3 are inversed (Figure 2F and 2G). This indicates that eIF4E3 replaces eIF4E1 in the eIF4F complex when the latter is sequestered by hypophosphorylated 4EBP1.

These results suggest that the PI3K/AKT/mTOR pathway regulates a function of eIF4E3, namely, its recruitment into an eIF4F complex. To confirm this, we examined the impact of upstream inhibitors of the mTOR pathway on the sedimentation profiles of eIF4E1, eIF4E3 and eIF4G1. We targeted PI3K activity using LY294002 (Hawkins et al., 2006; Vlahos et al., 1994). LY294002 caused a loss of AKT-Ser^473^, 4EBP1 and S6K1 phosphorylation (Figure 2H) mimicking Torin1 treatment (Figure 2I). The majority of eIF4E1 moved to the top of the gradient in fractions 8 and 9, and eIF4E3 was now found in the same fractions as eIF4G1. This confirms that eIF4E3 can assemble an eIF4F complex when the PI3K/AKT pathway is compromised. We then tried to mimic the effect in a physiological setting. The activation of mTORC1 is nutrient dependent with its Raptor and mLst8 subunits acting as nutrient sensors (Hara et al., 2002; Kim et al., 2002). We starved cells of nutrients and a carbon source by removing serum and glucose from the media. After two hours of starvation, AKT and S6K1 remained phosphorylated whereas 4EBP1 was hypophosphorylated (Figure 2J). A major part of eIF4E1 still co-sedimented with eIF4G1 (Figure 2K). However, some eIF4E3 was also detected in the eIF4G1 fraction (Fraction 4). Even though the effect was less marked, physiological conditions such as glucose starvation appear to promote the eIF4E3 interaction with eIF4G1.

### The eIF4E3 interactome

To validate that eIF4E3 could assemble a *bona fide* eIF4F complex we performed a yeast-2-hybrid (Y2H) screen to determine its partners. eIF4E3 was used as a prey and screened against a peptide library originating from human prostate cancer cell lines. This background was selected because eIF4E3 has been implicated in androgen-independent prostate cancer progression (Lin et al., 2007). About 50 million interactions were tested and 360 clones screened positive. Data mining revealed ten potential partners of eIF4E3 whose functions are listed in Table 1. A selected interaction domain (SID) for each prey sequence is depicted in Figure S2A. Among the ten partners were eIF4G1 and eIF4G3, consistent with previous observations that eIF4E3 could form an eIF4F(3) complex with eIF4G1 (Landon et al., 2014; Robert et al., 2020) and with eIF4G3 (Frydryskova et al., 2016). The eIF4G3 protein has been shown to associate with eIF4E2 during hypoxia (Ho et al., 2016; Uniacke et al., 2012), and interacts with eIF4E1 (Caron et al., 2004; Pyronnet et al., 2001).

**Table 1.**
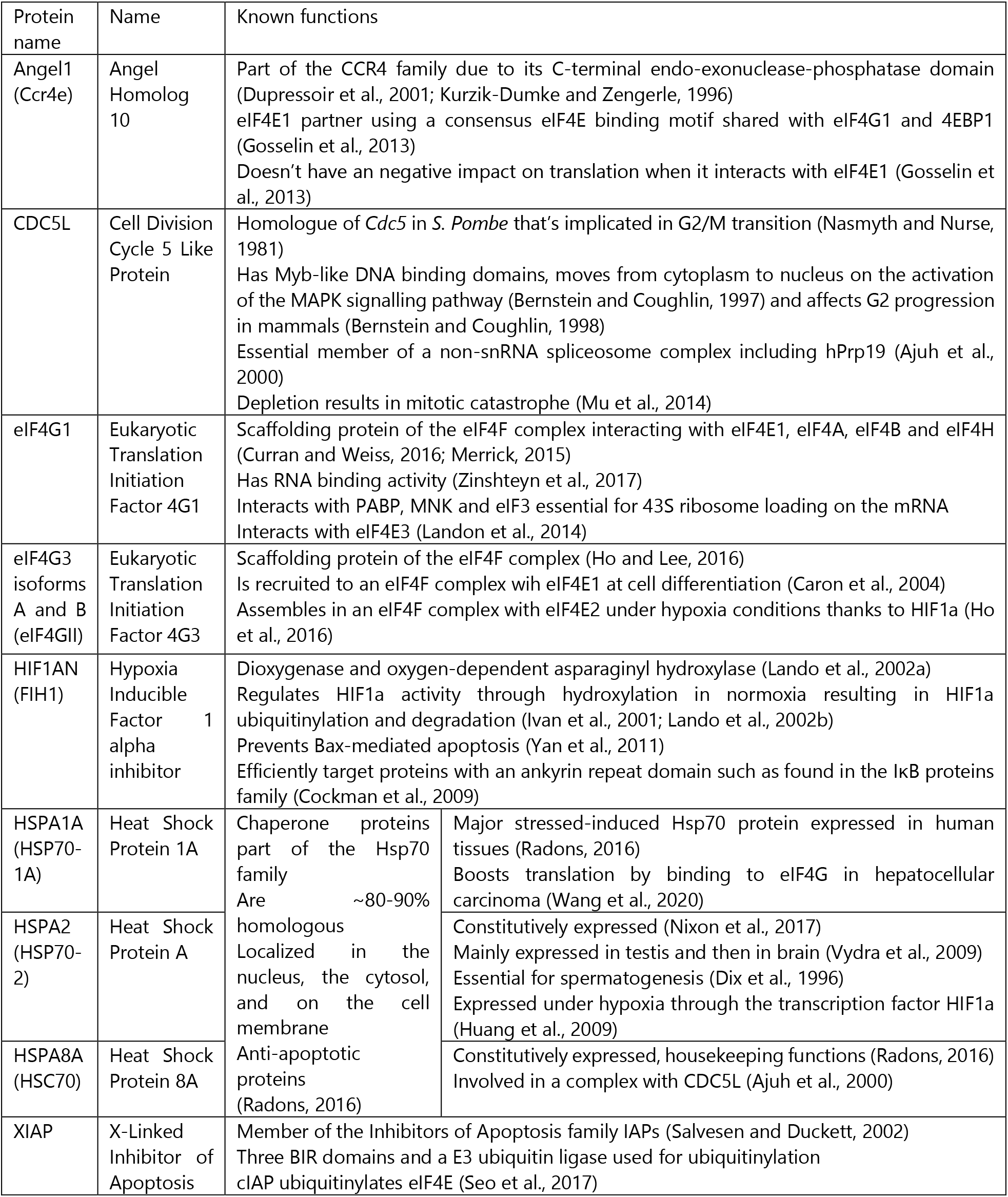
eIF4E3 partners as determined by Y2H assay and their known functions.

The SIDs on eIF4G1 and eIF4G3 were overlapping and corresponded to the reported binding site for eIF4E1 (Figure S2A), suggesting that eIF4E1 and eIF4E3 compete for the same surface on the scaffolding protein. To confirm the interaction, we performed co-IP. Cells transduced with human eIF4E3^HA^ were treated for two hours with Torin1 or vehicle, lysed and incubated with beads carrying covalently cross-linked HA antibody. The presence of eIF4G1 and eIF4G3 on the beads was then analysed by Western blot (Figure 3A). Both eIF4G1 and eIF4G3 co-IPed with eIF4E3^HA^ in Torin1 conditions confirming that eIF4E3 assembles into an eIF4F complex during stress (hereafter referred to as eIF4F^S^) with either eIF4G1 (eIF4F^S1^) or eIF4G3 (eIF4F^S3^). The seven other partners are not linked to translation and show no common motif with the exception of Angel1 with eIF4G1/3 (Figure S2B). Angel1 is a member of the CCR4 family (Dupressoir et al., 2001) and interacts with eIF4E1 via the YxxxxLΦ 4E binding motif. However, it plays no role in translational control (Gosselin et al., 2013). The other partners are CDC5L that is involved in the G2/M transition during the cell cycle; three heat shock proteins of the HSP70 family that share more than 80% homology (Radons, 2016); the HIF1α inhibitor HIF1AN and the X-linked inhibitor of Apoptosis XIAP. To confirm the interactome, we employed the Split Venus fragment technique that is a bimolecular fluorescent complementation assay (Ohashi and Mizuno, 2014). The C-term of eIF4E3 was fused to the N-term of Venus (VN210) and the putative partner protein to its C-term (VC210). Cells were co-transfected with eIF4E3-VN and either empty-VC or partner-VC. To test if the interaction responded to Torin1, cells were treated for two hours 2 days post-transfection. YFP was not observed with eIF4E3-VN alone or in the presence of empty-VC (Figure 3B). Even under these conditions of overexpression, eIF4G3-VC was observed in close proximity with eIF4E3-VN only in Torin1 treated cells, consistent with our co-IP results (Figure 3D). However, eIF4G1-VC, CDC5L-VC, HIF1AN-VC, HSPA8-VC and XIAP-VC gave signals independently of Torin1 treatment (compare Figure 3C and 3E-H and Figure S3). Interestingly, the signal localization was dependent on the partner tested. For eIF4G1-VC, eIF4G3-VC, HIF1AN-VC, HSPA8-VC and XIAP-VC, the signals were detected exclusively in the cytosol. CDC5L complementation aggregates were observed in the nucleus (Figure 3, top lane) with a staining reminiscent of nucleoli (Figure 3, bottom lane) (Yang et al., 2018). We observed cytosolic signals in cells entering mitosis (Figure 3, middle lane). Our results indicate that trace amounts of eIF4E3 may be present in the nucleus in association with CDC5L.

**Figure 3.**
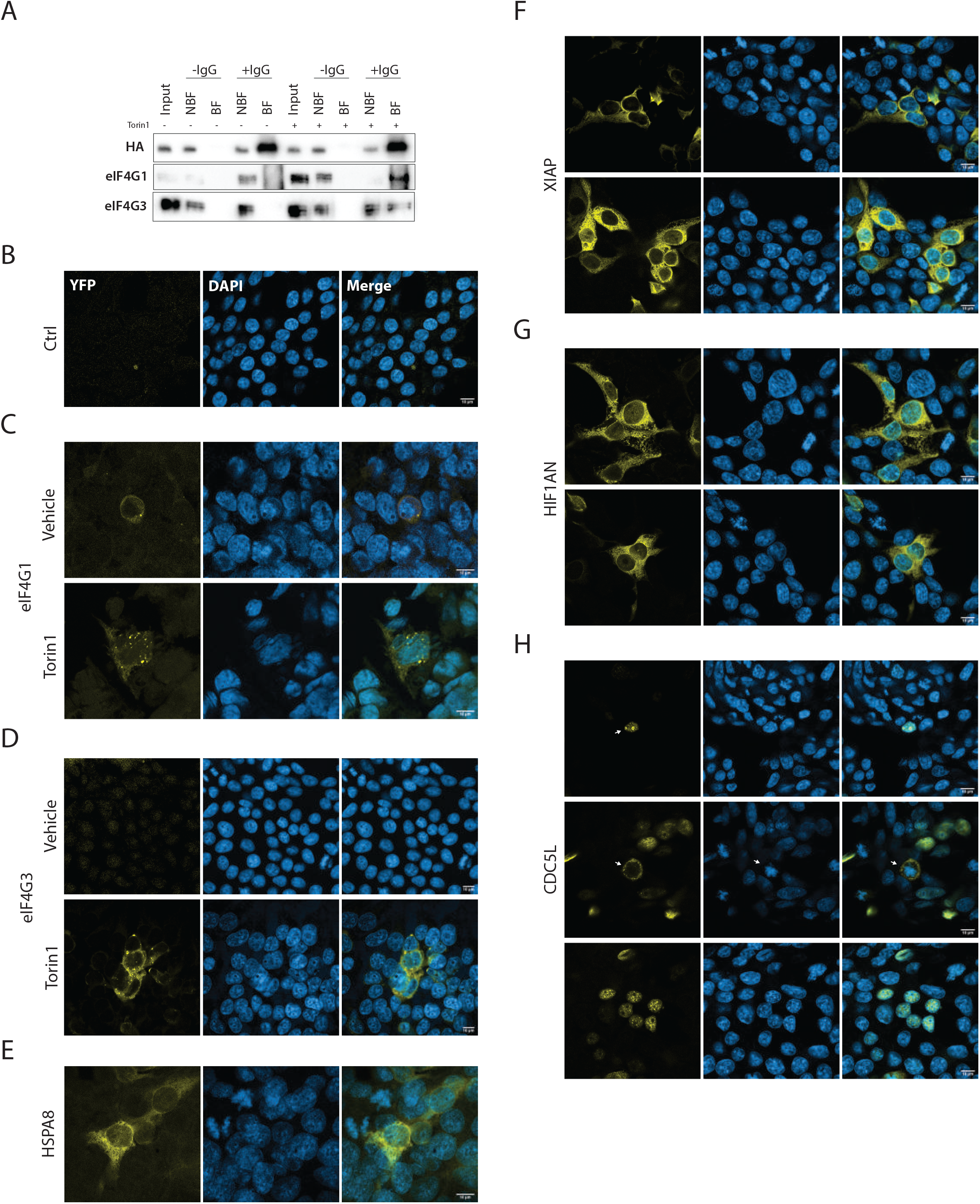
The eIF4E3 interactome. **A**. Western blot analysis of the co-IP assay using eIF4E3^HA^ and endogenous eIF4G1 or eIF4G3 starting from a lysate of HEK293T cells transduced with eIF4E3^HA^ treated with vehicle or with 250 nM Torin1 for 2 hours. Beads carrying covalently cross-linked Anti-HA Ab (+IgG) were used to immunoprecipitate eIF4E3^HA^ Beads without Ab served as a control (-IgG). **B-H**. Bimolecular complementation assay using Venus YFP in transfected HEK293T cells. Images were generated by confocal microscopy. Venus fragment 1 and Venus fragment 2 were fused to eIF4E3 and to one of the tested partners, respectively: empty vector (**B**), eIF4G1 (**C**), eIF4G3 (**D**), HSPA8 (**E**), XIAP (**F**), HIF1AN (**G**) and CDC5L (**H**).

### eIF4E3 impacts on the translatome in stress conditions: Polysome seeding based on 5’ TL length

With the confirmation that eIF4E3 forms an eIF4F^S^ complex, we sought to exploit a knockout (KO) approach to examine the impact of eIF4E3 loss on the translatome. As HEK293T cells express low levels of eIF4E3, we screened for a cell line in which protein expression was high (Figure S4). While eIF4E1 was found in all cell lines tested (Figure S4A), the expression level of eIF4E3 varied considerably (it was undetectable in the non-tumoural cell lines NIH3T3, MEF and MRC5, but highest in the mouse neuroblastoma cell line N2a) (Figure S4B). We therefore decided to pursue our analysis in N2a cells. Neuroblastoma cells represent cancer cells that arise from chromaffin cells mostly found in the medulla of the adrenal glands (Tsubota and Kadomatsu, 2017), a tissue in which eIF4E3 levels in the mouse are also high (https://www.mousephenotype.org/data/genes/MGI:1914142). We KOed eIF4E3 using CRISPR-Cas9 (Hsu et al., 2014). N2a cells were transfected with empty vector (control cells: ctrl) or with three gRNAs targeting exons 1, 2 and 4 (KO cells). Protein expression was reduced to undetectable levels (Figure 4A). We performed polysome profiling on ctrl and KO cells treated with vehicle or Torin1 for two hours. We did not observe marked changes in the profiles between ctrl and KO cells treated with vehicle (Figure 4B, first panel).

**Figure 4.**
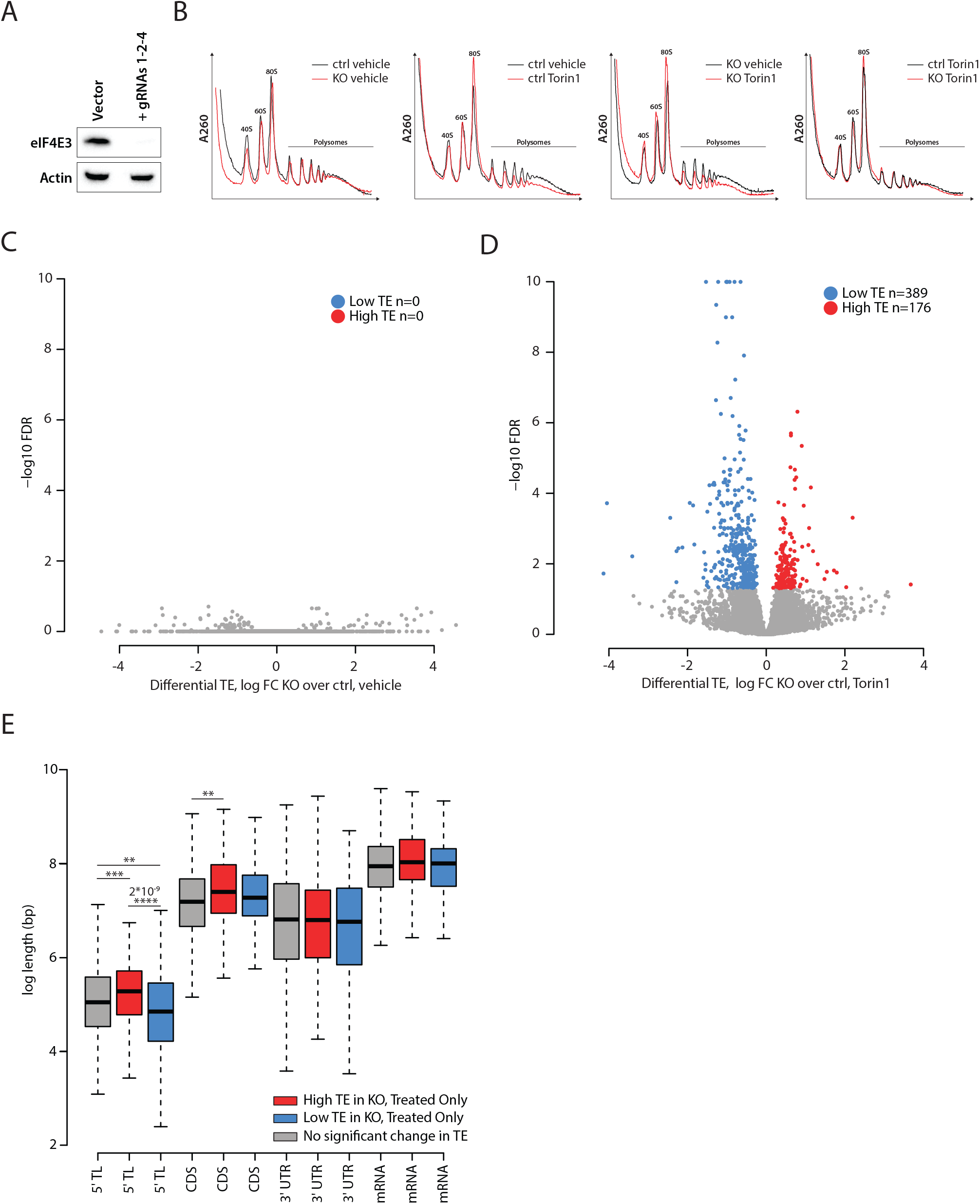
eIF4E3 is involved in translational re-programming during Torin1-induced stress. **A**. Western blot showing the level of endogenous eIF4E3 in N2a cells after transfection with empty vector or vector containing gRNAs targeting eIF4E3 following antibiotic selection. **B**. Polysome profiles of N2a cells control (ctrl) and eIF4E3 KO treated with vehicle or 250 nM Torin1 for 2 hours. **C-D**. Volcano plots of RiboDiff output, showing log2 fold change of differential translational efficiency (TE) comparing KO vs ctrl for vehicle (**C**) and Torin1 (**D**) against the resultant false discovery rate (FDR) of each comparison. Transcripts showing significantly higher or lower TE in KO are highlighted in red and blue, respectively. **E**. Boxplot comparing lengths of 5’TL, CDS, 3’UTR and mRNA for High TE, Low TE and No TE change groups. We note that 5’TL lengths are significantly higher and lower than control for high TE and low TE groups, respectively. High TE and Low TE groups were defined (using RiboDiff output) as transcripts with FDR<0.05, showing log2 TE fold changes for KO/ctrl that are positive and negative, respectively. The no change group was defined as having logFC < 0.05 and FDR>0.4 when comparing KO vs ctrl at the TE, ribo-seq and RNA-seq level. **: p-value < 0.01; ***: p-value < 0.001; p-value < 0.001. p-values < 0.0001 are indicated. No value indicates no significant changes.

However, Torin1 treatment reduced the heavy polysomal fraction in both (Figure 4B, second and third panels), but the effect was more pronounced in the KO (Figure 4B, fourth panel). To evaluate the impact of eIF4E3 on the translatome, we turned to ribosome profiling (ribo-seq) (Ingolia et al., 2009). This was performed on biological triplicates of ctrl and KO cells treated with vehicle or with Torin1 for 2 hours. The phosphorylation status of 4EBP1 was used to monitor drug treatment using an aliquot of the cell extract sampled before preparation of the ribo-seq library (Figure S5A). We obtained good reproducible data with the exception of an outlier in the N2a/vehicle ctrl that was discarded for the analysis (Figure S5B). The ribo-seq data showed clear triplicity throughout the annotated CDSs and correct footprint size distribution (Figure S5C and S5D). The transcriptome was also ascertained in all conditions (Figure S5E). This revealed that the steady state level of 36 mRNAs were significantly altered in the ctrl versus KO, with eIF4E3 being the most downregulated in the KO (Figure S6A). We also noted that Torin1 treatment modified the transcriptome (Figure S6A), an observation already reported (Park et al., 2017). The absence of eIF4E3 had a significant additional impact on the transcriptome with half of the differentially expressed (DE) genes (ctrl/Torin1 versus KO/Torin1) being downregulated and the other half upregulated (Figure S6A). To assess the impact of eIF4E3 on global translation, translation efficiency (TE) was evaluated from our ribo-seq and RNA-seq data. The TE was unchanged between ctrl and KO (Figure 4C). This is consistent with eIF4E3 playing no role in the translational readout in normal physiological conditions as it is not found in an eIF4F complex. However, Torin1 treatment had a major impact on the translatome in both cell lines, modifying the TE of a number of genes (Figure S6A). It is established that the translation of one mRNA subpopulation is highly mTOR dependent. These are referred to as the 5’ Terminal-Oligo-Pyrimidine (TOP) mRNAs and they share a common 5’ TL signature starting with a cytidine immediately after the 5’ cap followed by a 4-14 poly-pyrimidine stretch (Jefferies et al., 1994; Meyuhas, 2000; Thoreen et al., 2009). These mRNAs essentially code for ribosomal proteins and translation elongations factors (Iadevaia et al., 2008; Thoreen et al., 2009). TOP mRNAs selected from a list taken from (Thoreen et al., 2009) were downregulated in both cell lines after Torin1 treatment (Figure S6A, orange dots) confirming that the drug treatment worked. Their downregulation was similar in both cells lines, meaning that eIF4E3 does not impact TOP mRNA translation (Figure S6B, R = 0.96). When comparing the Torin1 treated cells, a considerable number of genes exhibited TE changes in the eIF4E3 KO (Figure 4D). The majority of these genes (ctrl/Torin1 versus KO/Torin1) had a lower TE, meaning their translation under the Torin1-induced stress is eIF4E3 dependent. Nevertheless, about one third had a higher TE, suggesting that eIF4E3 has an inhibitory effect on their translation in stress conditions. The ribosome occupancy on the CDS was similar in all four conditions and it did not change between the down and up regulated populations (Figure S6C-E). This result indicates that translation elongation was not altered in the four conditions; hence, TE changes reflect a defect at the level of initiation. As eIF4E3 is a cap binding protein, we looked for possible motif(s) or sequence signature(s) in the 5’ TL of the mRNA subpopulations responding to the KO under stress, namely, the neutral (no significant change in TE - ctrl/Torin1 versus KO/Torin1), stimulated (high TE) or inhibited (low TE). We were unable to find sequence motifs that correlated with each group. However, whereas the CDS, 3’ UTR and overall mRNA lengths did not vary within the three populations, 5’ TL length was significantly different (Figure 4E). The downregulated population had a significantly shorter 5’ TL compared to the neutral population, whereas the upregulated population had a significant longer 5’ TL (Figure 4E). This result suggests that eIF4E3 can impact both positively and negatively on the translational readout during stress and this correlates with 5’ TL length.

### N2a KO cells manifest defects in cellular metabolism under Torin1-induced stress

We were then interested to look at the impact of the KO on the cell phenotype under stress conditions. As the proteome is an excellent marker for the cellular phenotype and correlates better with the translatome than with the transcriptome (Smircich et al., 2015), we exploited only the ribo-seq data to determine what proteins are differentially expressed (Figure 5A). In agreement with the TE, the ribo-seq revealed that two-thirds of the mRNAs were downregulated (n = 492) and one-third upregulated (n = 278). KEGG analysis demonstrated that each population was enriched in defined pathways (Figure 5B and 5C) and in functions associated with different cellular compartments (Figure S7A and S7B).

**Figure 5.**
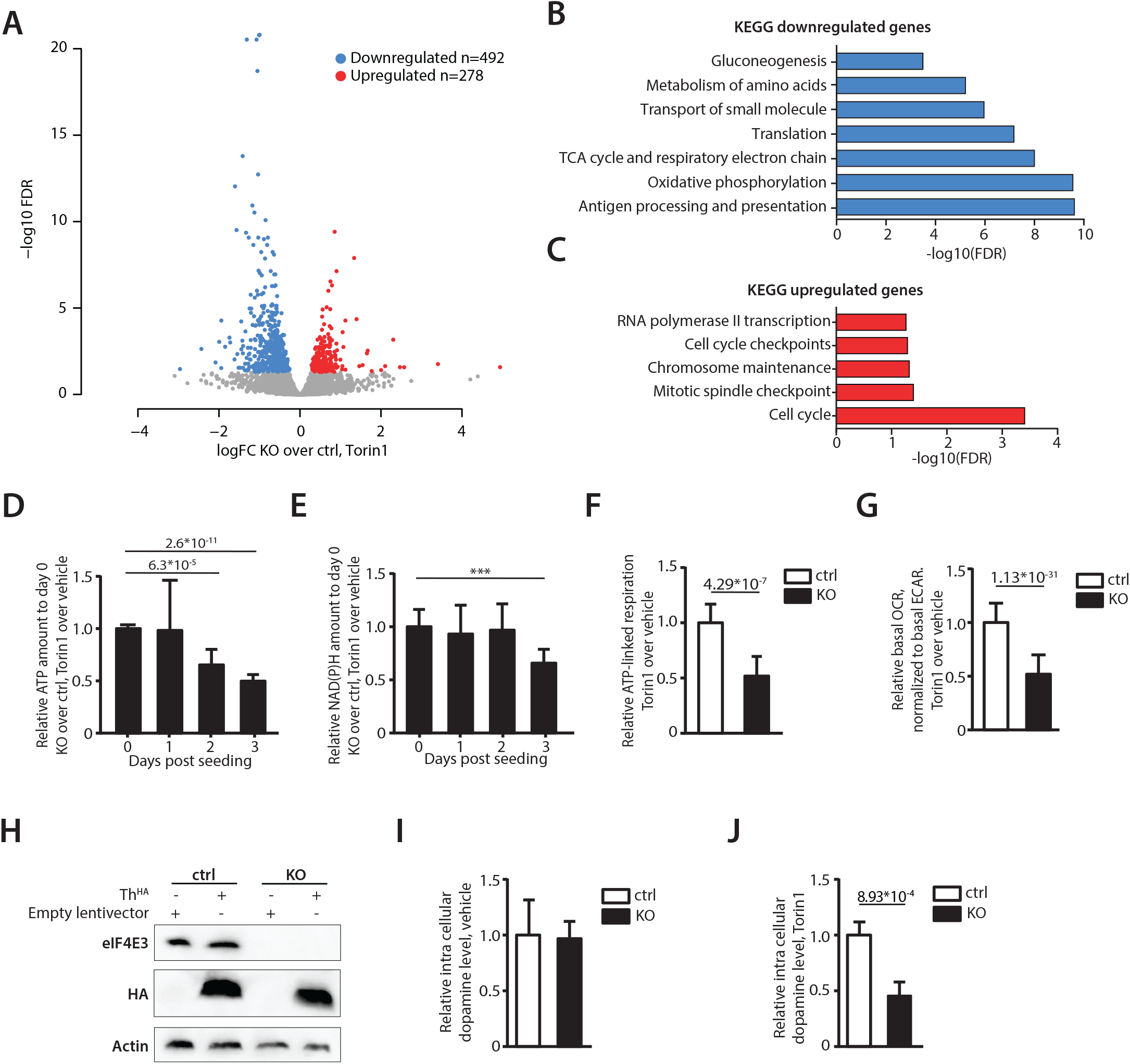
A role for eIF4E3 in the metabolic response during Torin1-induced stress. **A**. Volcano plot of edgeR output, showing log2 fold change of differential expression of ribosome profiling footprint (RPF) counts for each CDS, comparing KO and ctrl in Torin1. **B-C**. Barplot of FDR values for hypergeometric tests showing enrichment of various KEGG terms among genes downregulated (**B**) and upregulated (**C**) in KO vs ctrl, in Torin1. **D-E**. Histogram showing the relative ATP (**D**) or NAD(P)H (**E**) levels over time: Values represent the [KO treated]/[KO untreated] ratio relative to the [ctrl treated]/[ctrl untreated] ratio normalised to day 0. P-values are shown above. **F**. Relative ATP levels generated by cellular respiration in treated relative to untreated ctrl or KO cells. **G**. Relative oxygen consumption rate (OCR) relative to the extracellular acidification rate (ECAR) of ctrl or KO cells treated with Torin1 normalised to vehicle conditions as measured using the Mito Stress Test kit. **H**. Western blots using Abs for eIF4E3, HA and actin from N2a ctrl or KO cells transduced with empty lentivector or a vector expressing Th^HA^. **I**. Histogram of the dopamine levels in ctrl or KO cells treated with vehicle for 4 days. **J**. Histogram of the dopamine level in ctrl or KO cells treated with Torin1 for 4 days. **: p-value < 0.01; ***: p-value < 0.001; p-value < 0.001. p-values < 0.0001 were indicated directly. No value: no significant changes.

To validate the KEGG analysis we chose to explore two metabolic pathways. Firstly, to confirm that cellular metabolism was affected in the KO under stress, the levels of ATP and NAD(P)H were measured in ctrl and KO N2a cells. Since the impact of translatome changes on the proteome will also depend on protein stability, and mammalian proteins have half-lives ranging from hours to days, measurements were performed over several days of drug treatment. Cells were seeded and treated with vehicle or with Torin1 throughout the duration of the assay. Every 24 hours, ATP and NAD(P)H levels were assayed and their values normalised to day 0 and to the cells treated with vehicle. Both ATP and NAD(P)H levels showed a significant decrease at day 2 and 3 post-treatment (Figure 5D and 5E). To test if the reduction in intracellular ATP level was related to reduced mitochondrial respiration as suggested by the pathway enrichment analysis (Figure 5B), we probed the oxygen consumption rate using a 96-well metabolic analyser (Seahorse XF96, Agilent Technologies). Following three days of pre-treatment with vehicle or Torin1, ctrl and KO cells were seeded into 96 well plates and probed 36 hours later using a Mito Stress Test kit. The ATP linked respiration rate (difference between basal respiration and respiration following oligomycin addition), was modestly reduced in ctrl cells following Torin1 treatment (20% reduction), but markedly reduced in KO cells (60% reduction), confirming a reduction in ATP production through mitochondrial respiration in KO cells upon Torin1 treatment (Figure 5F). Similarly, KO cells displayed the lowest absolute basal respiration rate (oxygen consumption rate, OCR) as well as the lowest relative OCR when normalised to the basal glycolytic rate (extracellular acidification rate, ECAR). This relative respiration rate was 2-fold reduced in WT cells upon Torin1 treatment, but 3.4-fold reduced in KO cells (Figure 5G). We conclude that mitochondrial respiration is reduced in N2a cells upon Torin1 treatment and that this effect is markedly aggravated in the KO cell line, contributing to reduced ATP levels.

One of the hallmarks of cellular stress in neuroendocrine cells such as neuroblastoma cells is the production of catecholamine (Tsubota and Kadomatsu, 2017). However, N2a cells only produce dopamine when differentiated into dopaminergic cells. The differentiation process triggers the expression of tyrosine hydroxylase (Th) an enzyme that is normally weakly express in undifferentiated N2a cells (Tremblay et al., 2010). Th catalyses the conversion of L-Tyrosine to L-DOPA, an intermediate in dopamine synthesis. Mining of the ribo-seq data suggested that the KO under Torin1 conditions would also influence this metabolic pathway. To investigate this, both ctrl and KO cells were transduced with lentivectors carrying Th^HA^ (Figure 5H). This restored the production of dopamine in both backgrounds as evidenced by an increase in its intracellular (i.c.) levels (Figure S7C). Cells were then treated for four days with Torin1 or vehicle and i.c. dopamine levels determined. While we did not observe any significant change under vehicle conditions (Figure 5I), KO cells under Torin1 had about 50% less i.c. dopamine compared to control (Figure 5J).

## Discussion

The function of the eIF4E3 remains somewhat enigmatic. It has been described to function as a tumour suppressor via its ability to compete with eIF4E1 for the 5’ cap with an apparent inability to interact with eIF4G (Osborne et al., 2013), and as a promoter of tumourigenicity, a gain of function phenotype reminiscent of an oncogene (Abdelfattah et al., 2018; Lin et al., 2007). The KO mice are viable, and no predisposition to tumour formation was reported. To further “cloudy the waters”, studies concluded that eIF4E3 could interact with eIF4G1 and eIF4G3 (Joshi et al., 2004; Robert et al., 2020) but not the 4EBPs (Joshi et al., 2004) and its over-expression potently stimulated the expression of a reporter (Robert et al., 2020). Our own transient expression/co-IP assays are consistent with the body of these studies specifically with regards to eIF4E3s interaction with the 5’ cap plus eIF4G1 and its “non-interaction” with 4EBP (Figure 1). However, all these studies employed transient over-expression, *in-vitro* protein expression or purified tagged proteins/peptides. None actually probed what is happening with the endogenous protein in the cellular environment. Employing a sedimentation profiling protocol we observed that in normally growing cells, the endogenous eIF4E3 was not in an eIF4F complex, probably because it was unable to compete with the endogenous eIF4E1 for the eIF4G pool. However, upon Torin1 (an inhibitor of the mTOR kinase), or LY294002 (an inhibitor of PI3K) treatment, and the sequestering of eIF4E1 by the hypophosphorylated 4EBPs, the eIF4G pool is liberated and eIF4E3 can now assemble into an eIF4F complex (Figure 6). This observation is consistent with an eIF4G competition model and the observations of Landon and co-workers (Landon et al., 2014). Since all drug treatments mimic cellular stress pathways that target the PI3K/AKT/mTOR pathway we refer to this complex as eIF4F^S^ (Heberle et al., 2015; Koromilas, 2019; Reiling and Sabatini, 2006; Saxton and Sabatini, 2017). Furthermore, our interactome studies demonstrate that this complex will assemble around both eIF4G1 (eIF4F^S1^) and eIF4G3 (eIF4F^S3^). Roberts and co-workers, (Robert et al., 2020) recently proposed that up to eight eIF4F complexes could exist in the cell composed of different combinations of the eIF4E, eIF4G and eIF4A family members. These conclusions derived from an RNA tethering assay using over-expressed proteins. Our current study extends this observation by establishing that these alternative eIF4F complexes can form with endogenous proteins and their composition can be regulated by physiological changes in the cell, drug-targeted mTOR inhibition or even glucose starvation (Figure 6). Thus like eIF4E2, that assembles an eIF4F^H^ complex during hypoxia, eIF4E3 will assemble an eIF4F^S^ complex during stresses that target the availability of both eIF4E1 and eIF4E2 because both have been reported to interact with the 4EBPs (Joshi et al., 2004). It also confirms the model initially proposed by Ho and Lee, (Ho and Lee, 2016) in which they envisaged the assembly of specific eIF4F complexes as a response to changing physiological conditions, producing what they referred to as the “adaptive translatome”.

**Figure 6.**
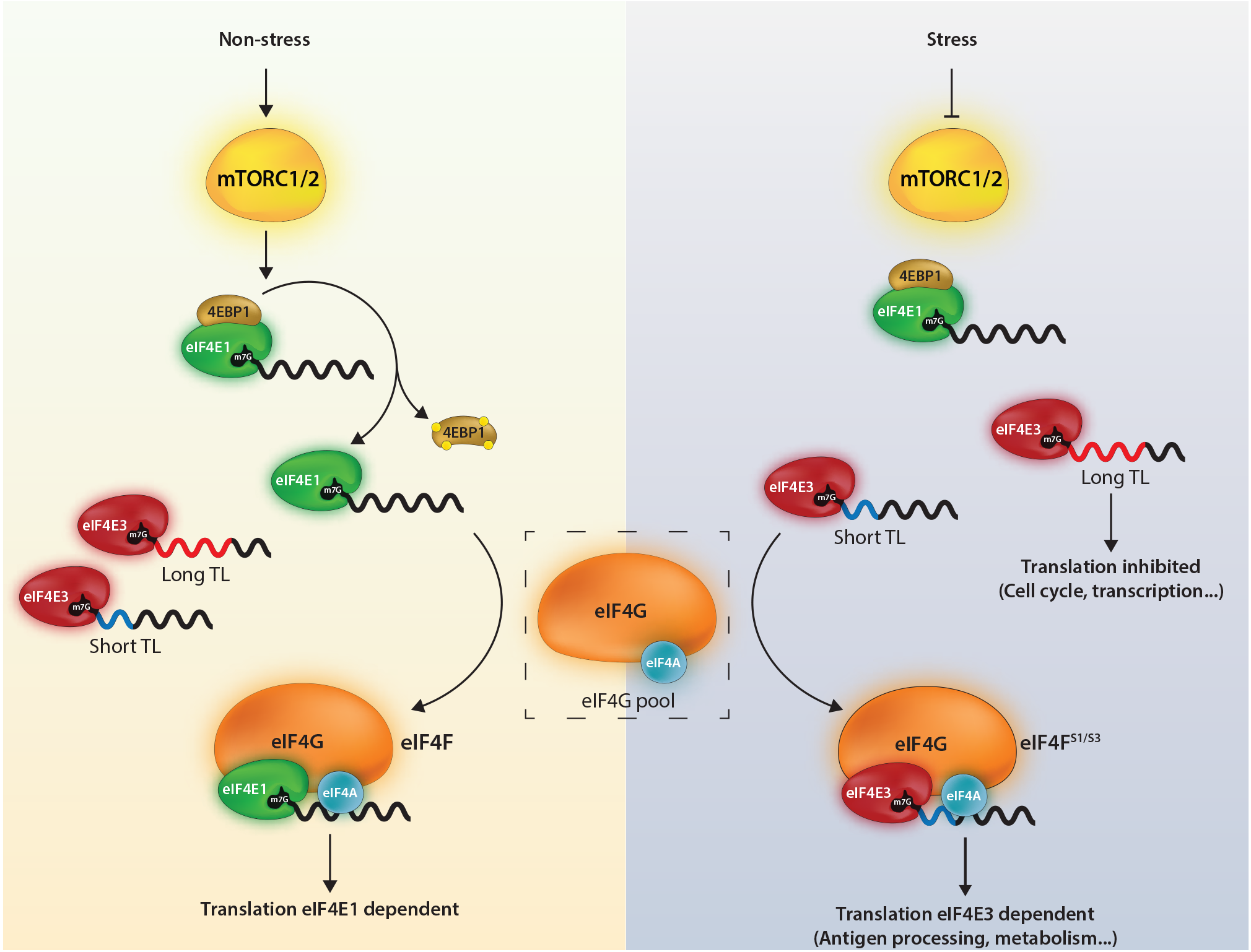
Model for the role of eIF4E3 in re-programming the translational readout during stress.

Having established a link between eIF4E3 and eIF4F^S^, what does this mean for the adaptive translatome? Studies that have previously probed this question used transient eIF4E3 over-expression in “non-stressed” cells analysed by polysomal profiling (Landon et al., 2014). However, ours is the first study on the comparative translatomes of a ctrl versus eIF4E3 KO using ribo-seq, a technique that gives a quantitative “snap-shot” of the translatome, after 2 hrs of Torin1 treatment (Andreev et al., 2015; Ingolia et al., 2009). It reveals that the impact of eIF4E3 loss only becomes apparent upon stress, a result compatible with the inability to assemble an eIF4F^S^ complex in the KO background. The translational response to stress is complexed, but it turns around two key initiation factors, namely eIF2 and eIF4E (Advani and Ivanov, 2019; Koromilas, 2019; Reiling and Sabatini, 2006). As alluded to in the introduction, despite the fact that the ISR reduces global protein expression, mRNA subpopulations continue to be expressed. This involves a selective re-seeding of the polysome with mRNA transcripts carrying uORFs within their 5’TL (Advani and Ivanov, 2019; Andreev et al., 2015). Decoding of the CDS is then assured by a mechanism referred to as translational reinitiation (Gunišová et al., 2018; Peabody and Berg, 1986; Skabkin et al., 2013). Hence the adaptive modification of the translatome is responding to features within the mRNA 5’ TL. Torin1 induced translational stress targets mainly the availability of eIF4E1 via eIF4E1-4EBP1 (Heberle et al., 2015; Jin et al., 2020). Our KO studies reveal that it is specifically as a response to this stress that eIF4E3 drives the formation of the adaptive translatome. In the absence of the drug, the translatomes of ctrl and KO are very similar (compare Figure 4C and 4D). As with the ISR, this re-seeding phenotype correlated with a specific feature within the 5’ TL, in this case length (Figure 4E). Transcripts with low TE in the KO/Torin1 cells tended to have shorter 5’ TLs and those with high TE longer TLs, relative to the non-responsive mRNA population. The high TE population also exhibited a G/C distribution downstream of the 5’ cap significantly different from both the low TE and non-responsive populations (Figure S7D). Therefore, under stress, the eIF4F^S^ complex will preferentially engage and drive the expression of transcripts with short 5’ TLs (Figure 6). This is somewhat reminiscent of the behaviour of eIF4E1, which also tends to select short 5’ TLs (Koromilas et al., 1992). Many housekeeping genes carry short 5’ TLs or TISU (translation initiator of short 5′ TL) elements (Elfakess and Dikstein, 2008; Elfakess et al., 2011; Sinvani et al., 2015). These genes tend to regulate metabolic pathways essential for cell survival. The results of our KEGG analysis of the downregulated gene set would suggest that the eIF4F^S^-driven expression serves to ensure that at least basal levels of metabolic function are maintained (Figure 5B). We experimentally confirmed this by demonstrating that mitochondrial respiration was negatively impacted by the loss of eIF4E3 upon extended Torin1-induced stress (Figure 5D-G). What is happening at the molecular level with regards to how the eIF4F^S^ selects these short TLs is unclear. Recent studies position the eIF4E1-4EBP on the 5’ cap (Jin et al., 2020; Ptushkina et al., 1999). The eIF4E1 has higher cap affinity than eIF4E3 and would presumably be positioned on the most accessible 5’ ends, i.e. those of short non-structured 5’ TLs. However, one must assume that during stress, the eIF4F^S^ complex can displace the eIF4E1-4EBP to promote PIC recruitment or that eIF4E3 already sits on the 5’ cap of specific transcript subpopulations, possibly because of an interaction with additional RNA binding proteins (although none were signalled on the Y2H analysis), awaiting the release of eIF4G. Curiously, we also observe a significant group of genes whose ribosome occupancy increased in the KO during Torin1 treatment. Modelling this effect is by no means evident, as it would seem that eIF4E3 is acting as a translational repressor on mRNA subpopulation(s). These are expressed only in the absence of eIF4E3 and only during stress. Aside from the effect on “house-keeping” functions, we also probed an N2a cell-specific function that appeared during our data mining. We were also able to demonstrate changes in intracellular dopamine levels in ctrl versus KO following extended Torin1 treatment (Figure 5I and 5J). Long drug treatment almost certainly introduces secondary effects, particularly in the transcriptional program. However, it takes time for changes in the translatome to impact the proteome because many of these metabolic enzymes are stable (Schwanhäusser et al., 2011).

The Y2H interactome study also produced a number of hits for proteins that are not thought to play any direct role in translational control (Table 1). Some of these we confirmed experimentally but unlike the interaction with the eIF4Gs, these occurred independently of stress. This included the transcription factor CDC5L, essential for the G2/M transition (Mu et al., 2014) and involved in the spliceosome complex (Ajuh et al., 2000). Interestingly, Abdelfattah and coworkers (Abdelfattah et al., 2018) published that eIF4E3 is required for G2/M progression in medulloblastoma cells. CDC5L shuttles from the cytoplasm to the nucleus upon activation of the MAPK pathway (Ajuh et al., 2000). eIF4E3 is essentially a cytoplasmic protein although it has been observed in the nucleus upon over-expression (Osborne et al., 2013). eIF4E1 is also known to shuttle between nuclear and cytoplasmic compartments via the transporter eIF4ENIF1 (Dostie et al., 2000). However, this transporter was not flagged in our interactome study. CDC5L may therefore be serving as the eIF4E3 transporter linking eIF4E3 function to the cell cycle.

These results suggest that eIF4E3 has “off-site” and possibly “moon-lighting” functions (Jeffery, 1999) i.e. functions outside the formation of the eIF4F^S^ complex. Such activities already exist for certain initiation factors (Wolf et al., 2020) and ribosomal proteins (Warner and McIntosh, 2009). These would occur in the absence of cellular stress exploiting the “free-pool” of eIF4E3 and would be non-essential.

### Conclusion

Bi-directional cross talk between the mTOR and ISR pathways has been extensively documented (Appenzeller-Herzog and Hall, 2012; Gandin et al., 2016; Nikonorova et al., 2018; Wengrod and Gardner, 2015; Zhang et al., 2018). Indeed, a recent translatome/proteome analysis of the cellular stress response concluded that all stress pathways converged with the aim of conserving a protein readout that assures cell function and survival (Klann et al., 2020). We propose that eIF4E3 serves as a second arm of the ISR responding to changes in the PI3K/AKT/mTOR pathway that compromise eIF4E1-mediated ribosome recruitment. The formation of eIF4F^S^ serves to ensure continued cap-dependent PIC recruitment and a selective re-seeding of the polysome (Figure 6). It provides a molecular model for how mTOR down-regulation can increase resistance to some types of stress (Reiling and Sabatini, 2006).

## Acknowledgments

This work was supported by the University of Geneva, the Swiss Science Foundation (31003A_175560) the Société Académique de Genève, the Ernst and Lucie Schmidheiny Foundation, the Ligue Genevoise Contre le Cancer and the Fondation Pour la Lutte Contre le Cancer. JK is funded from the European Research Council (ERC) under the European Union’s Horizon 2020 research and innovation programme under Grant agreement no. 695596.

## Conceptualisation

B.W. and J.A.C.

## Methodology

B.W. and J.A.C.

## Formal Analysis

G.E.A.

## Investigation

B.W., J.K., K.A., P.J.-G.

## Writing – Original Draft

B.W. and J.A.C. Supervision: J.A.C.

## Project Administration

B.W. and J.A.C.

## Funding Acquisition

J.A.C.

## Material and Methods

### Cell culture and treatment

HEK293T, N2a, NIH 3T3 and MEF cells were grown at 37°C in a humidified 5% CO_2_ chamber. The cells were cultured in Dulbecco’s Modified Eagle Medium (DMEM) supplemented by 1% penicillin/streptomycin (Gibco) and 10% fœtal bovine serum (Gibco) for HEK293T, N2a and MEF cells or 5% fœtal bovine serum for NIH 3T3 cells. The DMEM used for N2a cells was pyruvate free.

#### Drug Treatment

Cells were treated for 2 hrs with vehicle (DMSO) or with 250 nM Torin1 (Tocris Biosciences), or for 1 hr with 50 μM Ly29004 (Tocris Biosciences) before collection. For glucose starvation, HEK293T cells were washed with PBS and incubated with glucose-free DMEM (Gibco) supplemented with dialysed serum and pyruvate for 2 hrs prior to harvesting.

Protein stability was determined by treating cells with 100 μg/mL of cycloheximide for the times indicated in the text. Torin1 was kept in the media throughout the treatment.

#### N2a eIF4E3 KO

To generate N2a control and eIF4E3 KO cells, N2a cells were transfected with pSpCas9(BB)-Puro empty vector or vectors expressing each of the three gRNAs targeting the eIF4E3 exons (meIF4E3 gRNA exon 1, exon 2, exon 4) using Lipofectamine 2000 (Invitrogen). pSpCas9(BB)-Puro was a generous gift from Dr. Rabi Murr (University of Geneva, Switzerland). Cells were selected by treatment with puromycin which was added 72 hours post-transfection and maintained for 10 days. Surviving cells were then processed for a second round of transfection and selection.

### Viral particles production and transduction

HEK293T cells at 80% confluence were transfected with the second-generation packaging and envelope vectors (pWPI: https://www.addgene.org/12254/). Viral particles, collected and filtered through a 0.45 μm filter at 48 hrs post-transfection, were used to transduce HEK293T and N2a cells. Five days later, GFP expressing cells were sorted and selected using 3 μM puromycin (Sigma).

### Western blot

Protein extracts prepared in Laemmli buffer (50 mM Tris HCl pH 6.8, 8% (v/v) glycerol, 4% β-mercapto-ethanol, 2% SDS, 0.015% Bromophenol blue) were resolved on SDS-polyacrylamide gels and electro-transferred to PVDF. Antibodies used in this study were: anti-eIF4E1 (Cell Signaling, #9742S), anti-eIF4E3 (Protintech, #17282-1-AP), anti-eIF4G1 (Santa Cruz, sc-133155), anti-eIF4G3 (ThermoScientific PA5-31101), sc-133155), anti-eIF2α (Invitrogen, #44728G), anti-HA (Covance clone 16B12), anti-FLAG (M2 antibody, Sigma), anti-4EBP1 (Cell Signalling, #9452), anti-actin (Millipore, #MAB1501), anti-RPS6 (Cell Signalling, #2317), anti-Phospho-Akt (Ser473) (Cell Signaling, #9271), anti-AKT (Cell Signaling, #9272), anti-phospho S6 kinase (Thr389) (Cell Signalling, #9205), anti-p70 RPS6 kinase (Cell Signaling, #9202), goat anti-mouse HRP secondary antibody (Bio-Rad) and goat anti-rabbit HRP secondary antibody (Bio-Rad). The Anti-MYC tag was a gift from Prof. Dominique Soldati (University of Geneva, Switzerland) and anti-eIF4A was a gift from Prof. Michael Altmann (University of Bern, Switzerland). Immunoblots were quantitated using Image Lab (Biorad).

### Cap pull down

HEK293T cells were transfected with wild type and tryptophan mutants of HA-tagged eIF4E1 and eIF4E3 using calcium phosphate. Cells were lysed in cap binding buffer (50 mM Tris HCl pH 7.6, 150 mM NaCl, 2 mM EDTA, 1% (v/v) Triton, 0.5% (v/v) NP40). 500 μg of protein was incubated overnight at 4°C with m7GTP agarose beads (Jena biosciences, #AC-155S). Beads were washed with cap binding buffer and suspended in 20 μL Laemmli buffer. Input, non-binding and binding fractions were resolved on a polyacrylamide gel.

### Polysome profiling

Polysome profiling was performed as previously described (Dieudonne et al., 2015). Briefly, 20-60% sucrose (Sigma) gradients were prepared manually in 100 mM KCl, 0.2 mM MgCl_2_, 20 mM HEPES and 2 mM DTT. N2a cells were treated for 5 minutes with 50 μg/mL of cycloheximide (Sigma) and then collected in cold PBS containing 100 μg/mL cycloheximide. Cells were pelleted and lysed in polysome lysis buffer (100 mM KCl, 50 mM Tris-HCL pH 7.6, 1.5 mM MgCl_2_, 1 mM DTT, 1 mg/mL Heparin (Sigma), 1.5% (v/v) NP-40, 100 μg/mL cycloheximide) supplemented with protease cocktail inhibitor EDTA-free (Roche) on ice for 20 mins. Lysates were cleared by centrifugation (14000g) and supernatants were loaded onto the gradients. These were centrifuged in a SW41 rotor for three and a half hours at 35000 rpm at 4 °C. After centrifugation, the gradients were analysed through an UV-lamp and an Absorbance detector while being collected in 1 mL fractions using a Foxi Junior Fraction Collector (Isco).

### RNA-seq

N2a cell total RNA was purified using the TRIzol Reagent following the manufacturers’ instructions. Libraries were prepared with the TruSeq stranded RNA Library Prep (Illumina) and sequenced at the iGE3 Genomic Platform (UNIGE) on a Hiseq 4000.

### Ribosome profiling

N2a cells were treated for 2 hours with vehicle or Torin1 and then for 5 minutes with 100 μg/mL of cycloheximide (Sigma). Cells were collected in cold PBS supplemented with 100 μg/mL of cycloheximide and then lysed on ice for 15 min in mammalian lysis buffer (100 mM KCl, 50 mM HEPES pH 7, 1.5 mM MgCl_2_, 1 mM DTT, 1% (v/v) Triton, 5U DNase I (Ambion), 100 μg/mL cycloheximide) supplemented with protease cocktail inhibitor EDTA-free (Roche). Lysates were clarified by centrifugation (10000g) and then digested with 7.5U of RNase I (Ambion). RNA was pelleted and purified on a 1 M sucrose cushion by centrifugation in a S45A rotor at 40000 rpm for 4 hours at 4 °C. Pellets were resuspended in TRIzol Reagent (Invitrogen) and RNA was extracted according to the manufacturers instructions. Ribosomal RNA was depleted from the extract (RiboMinus v2 Eukaryote Kit, Invitrogen). RNA fragments with a size ranging from 28 to 34 nt were extracted from a polyacrylamide urea gel and purified by precipitation. RNA was dephosphorylated using T4 Polynucleotide Kinase (NEB). The reverse transcriptase linker was then ligated with T4 RNA Ligase 2 truncated (NEB) and reverse transcribed using M-MLV RNase H minus (Promega). cDNA products were purified on gel and amplified by PCR using the Phusion polymerase (NEB). Libraries were sequenced at the iGE3 Genomic Platform (UNIGE) on a Hiseq 4000.

### RiboSeq and RNASeq Mapping

For the Ribo-Seq samples, all fastq files were adaptor stripped using cutadapt. Only trimmed reads were retained, with a minimum length of 15 and a quality cutoff of 2 (parameters: -a CTGTAGGCACCATCAAT -- trimmed-only --minimum-length = 15 --quality-cutoff = 2). Histograms were produced of ribosome footprint lengths and reads were retained if the trimmed size was between 25 and 35. For all Ribo-Seq and RNA-Seq samples, reads were mapped, using default parameters, with HISAT2 (Kim et al., 2015), using their pre-prepared UCSC mm10 genome and index. Only primary alignments were retained and reads were removed if they mapped to rRNA, tRNA and pseudogenes according to mm10 RepeatMasker definitions from UCSC. A full set of transcript and CDS sequences for Ensembl release 84 was then established. Only canonical transcripts [defined by mm10 knownCanonical table, downloaded from UCSC] were retained with their corresponding CDS. Reads were then mapped to the canonical transcriptome with bowtie2 (Langmead and Salzberg, 2012) using default parameters.

### Ribo-Seq Analysis

The P-site position of each read was predicted by riboWaltz (Lauria et al., 2018) and confirmed by inspection. Counts were made by aggregating P-sites overlapping with the CDS and P-sites Per Kilobase Million (PPKMs) were then generated through normalising by CDS length and total counts for the sample. Differential expression was performed pairwise between Ctrl or KO triplicates in the presence of Torin1 or Vehicle using edgeR (Robinson et al., 2010) on default settings. Transcripts were only kept in the analysis if they had a CPM > 1 in all triplicates for either at least one of the conditions in the pairwise comparison. For further analysis transcripts were filtered if their CDS length was not a multiple of three and if they did not begin with a standard start codon (Lawrence et al., 2000) and end with a standard stop codon (UAG, UGA, UAA). This left 20351 transcripts. Scaled plots summarizing the p-site depth profile over all relevant genes for the whole CDS were plotted by splitting every CDS in the gene group into 100 equal bins and aggregating the number of p-sites falling in each. They were normalised by counts for each CDS and total counts genome-wide.

### RNA-Seq Analysis

Counts were made by aggregating any reads overlapping with the CDS and RPKMs were then generated through normalising by CDS length and total counts for the sample. Differential expression was performed as with the Ribo-Seq.

### Translational Efficiency (TE) Analysis

TE was assessed using RiboDiff (Zhong et al., 2017) with default parameters with the same Ribo-Seq and RNA-Seq samples as input, using the same expression pre-filters as the edgeR differential expression analysis.

### Bimolecular Fluorescent Complementation assay

Yeast vectors containing Venus fragments were a generous gift from Prof. Martine Collart (University of Geneva, Switzerland). The Venus fragments were transferred into a pcDNA3 backbone. EIF4E3 was fused upstream of Venus fragment 1 at the SalI site. The partner was fused to the NotI site upstream of Venus Fragment 2. Details of the cloning strategy are available in the snapgene file. HEK293T cells were transfected with both plasmids using calcium phosphate at 24 hours post-seeding. At 48 hours post transfection, cells were treated with vehicle or Torin1 for 2 hours. Cells were fixed in methanol at −20°C, stained by DAPI (Sigma) and analysed by confocal microscopy (Zeiss).

### ATP and NADH assay

5.10^4^ control and eIF4E3 KO N2a cells were seeded into a 96 well dish and treated with vehicle or Torin1 for the duration of the assay. At 4 hours post seeding and then every subsequent 24 hours, ATP and NADH were independently measured using the Cell Glo Titer (Promega) and MTT reagent (Promega) following the manufacturers instructions.

### Seahorse analysis

N2a ctrl or KO cells were pretreated for three days with vehicle or Torin1. 3.10^4^ cells were seeded into 96 well plates suitable for a 96-well metabolic analyser (Seahorse XF96, Agilent Technologies). 24 hours later and prior to the assay, cells were washed once with Seahorse XF Base Medium (Agilent Technologies) and incubated with 180 μl of Seahorse XF Base Medium (Agilent Technologies) supplemented with 10 mM glucose and 2 mM glutamine. Cell metabolism was probed using a Mito Stress Test kit (Agilent Technologies). The measurement cycles (mix, wait and measure) were performed according to the standard settings. Three measurements of basal respiration were performed followed by three measurement cycles after the serial addition of 2 μM oligomycin, 0.5 μM carbonyl cyanide-p-trifluoromethoxyphenylhydrazone (FCCP) and 0.5 μM rotenone and antimycin A. The experiment was repeated twice, each time measuring more than 20 technical replicates. The ATP linked respiration rate, basal respiration rate and extracellular acidification rate were analysed using Wave (Agilent Technologies).

### Dopamine quantification

N2a ctrl and KO cells were plated in 6 well plates and treated with vehicle or with Torin1 for the duration of the assay. Four days after treatment, cells were collected in cold PBS, lysed in 0.1% (v/v) Triton X-100 and sonicated using a Branson Sonifier 450 (Branson, Danbury, CT, USA) at full power for 30 seconds. Cell extracts were then centrifuged at 3’000 g for 1 min and the supernatant was used to measure intracellular dopamine through extraction in activated alumina and quantified by ultraperformance liquid chromatography-tandem mass spectrometry (UPLC-MS/MS) (Dunand et al., 2013). Values were normalised to the protein amounts.

### Glycerol gradients

Glycerol gradients were prepared as previously described (Legrand et al., 2015). HEK293T cells were lysed in polysome lysis buffer without cycloheximide. Cell extracts were loaded onto a 5-20% linear glycerol gradient in 100 mM KCl, 5 mM MgCl_2_, and 20 mM HEPES prepared in an SW60 tube. Gradients were centrifuged for 22 hours at 40000 rpm in a SW60 rotor at 4 °C. After centrifugation, 10 fractions of 400 μL were collected from the bottom of the tube. Proteins were precipitated by methanol/chloroform precipitation and analysed by Western blotting. The pellet was resuspended directly in 40 μL of Laemmli buffer.

### Yeast-two-hybrid

Y2H was performed by Hybrigenics Services (https://www.hybrigenics-services.com/) against the Human Cancer Prostate_RP1 library.

### Co-immunoprecipitation

#### FLAG pull down

HEK293T cells were co-transfected with ^MYC-HIS^4EBP1 (kindly provided by Prof. Chris Proud, South Australian Health and Medical Research Institute) and either human eIF4E3^FLAG^ or murine eIF4E1^FLAG^. They were lysed in polysome lysis buffer containing 100 μg/mL cycloheximide and protease cocktail inhibitor EDTA-free (Roche)). 750 μgs of protein were incubated with 20 μL of FLAG-beads (Roche) overnight at 4°C. Beads were washed X3 in polysome lysis buffer, resuspended in Laemmli sample buffer and analysed on a SDS-polyacrylamide gel.

#### Histidine pull down

HEK293T cells were co-transfected with ^MYC-HIS^4EBP1 and either human eIF4E3^HA^ or murine eIF4E3^HA^. Cells were lysed in Talon IP buffer containing 100 mM NaCl, 0.5% (v/v) NP40, 40 mM Tris-HCl pH 7.6, 150 mM KCl and 2 mM MgCl_2_. 750 μgs of protein were incubated with 25 μL of TALON Metal Affinity Bead suspension (BD Biosciences) in a 500 μL final volume for 30 min at 4°C with slow rotation. Beads were recovered and washed three times with 500 μL Talon IP buffer. Proteins were eluted in 200 μM imidazole and resolved on a SDS polyacrylamide gel.

#### Co-Immunoprecipitation

HEK293T transduced with eIF4E3^HA^ and treated or non-treaated with Torin1 were collected in cold PBS before being lysed in IP buffer (50 mM HEPES, 2 mM EDTA, 10 mM Sodium Pyrophosphate, 10 mM b-Glycerophosphate, 40 mM NaCl, 1% (v/v) Triton X-100) on ice. Protein G magnetic beads were incubated with anti-HA antibody before being cross-linked with DSS (Thermo Scientific #88805). They were then incubated with cell lysate overnight at 4 °C, gently washed in IP buffer 2 times and finally resuspended in Laemmli sample buffer and resolved on a SDS polyacrylamide gel.

### DNA cloning

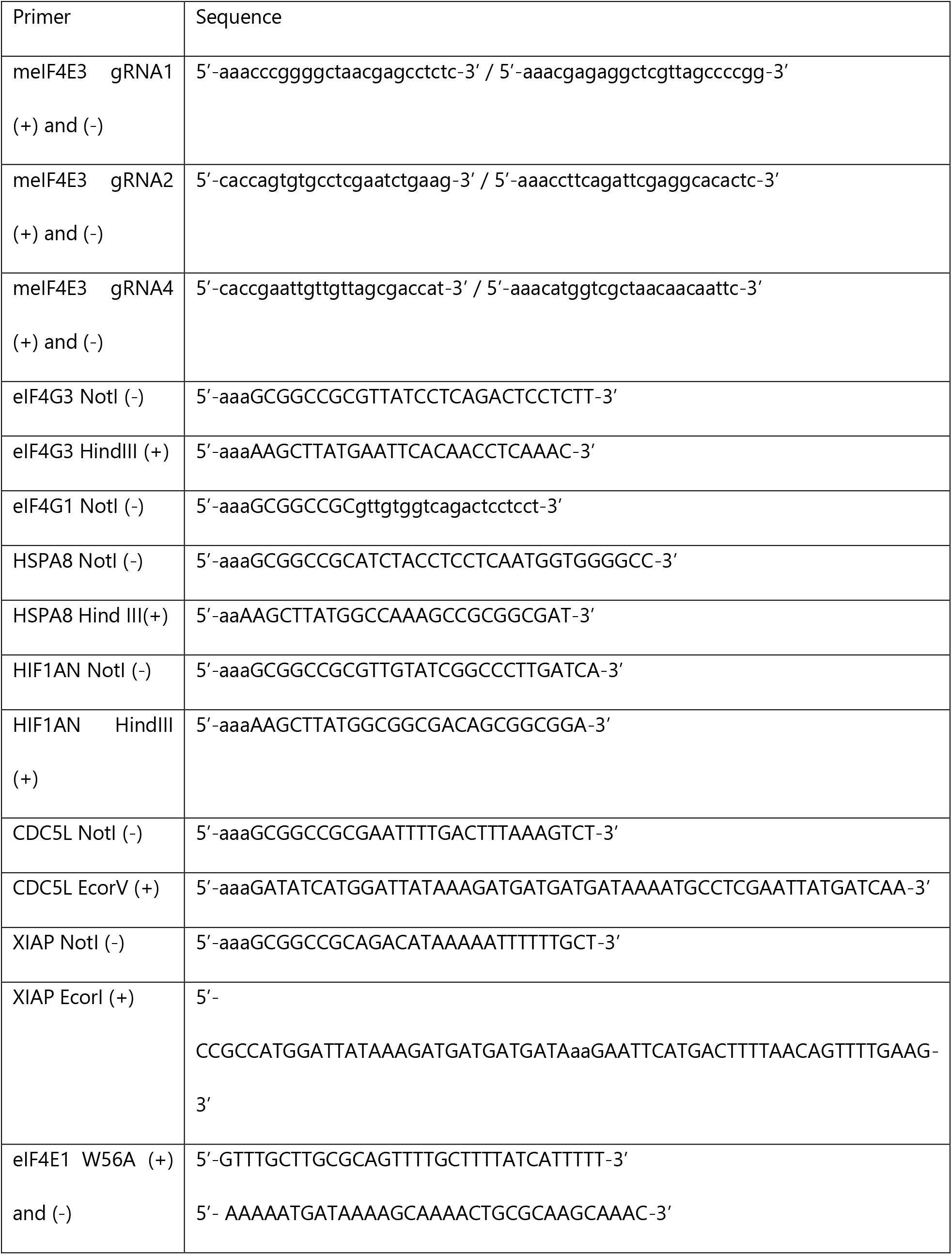
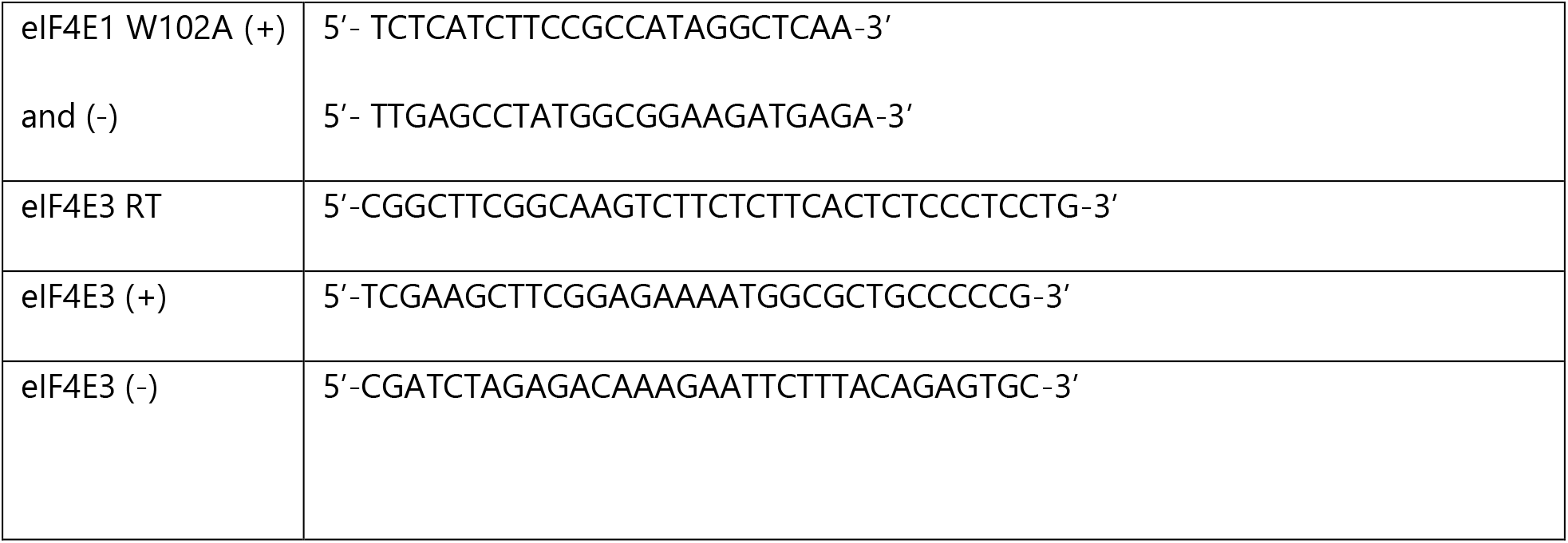

The pcDNA3 HA eIF4GI (1-1599) plasmid was a gift from Prof. Nahum Sonenberg (http://n2t.net/addgene) (Yanagiya et al., 2009). The pcDNA5/FRT/TO V5 HSPA8 plasmid was a gift from Prof. Harm Kampinga (http://n2t.net/addgene:19514) (Hageman and Kampinga, 2009). eIF4G3 (Biocat #BC094683-seq-TCHS1003-GVO-TRI) and Th (Biocat #BC156668-seq-TOMS6004-GVO-TRI) were purchased from BioCat. The pcDNA3 eIF4E1-FLAG S53A clone was a generous gift from Prof. David Sabatini (USA).

---

**Figure S1.**
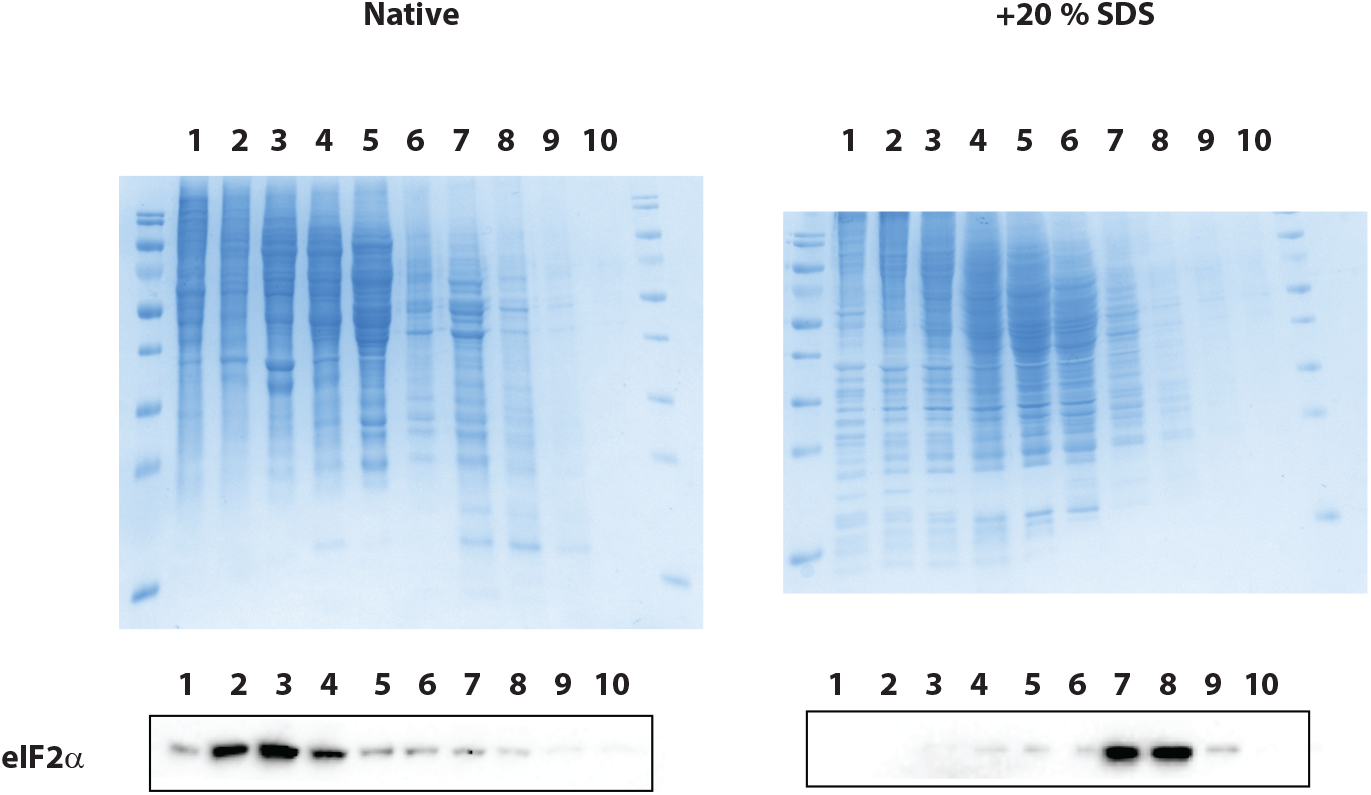
Upper pannel: Coomassie stained gel of the glycerol gradient sedimentation profiles from HEK293T cell lysates prepared under native and denatured (20% SDS treatment prior to gradient loading) conditions. Lower pannel: Western blot showing the sedimentation profile of eIF2a in the native and deternatured (+20% SDS) lysates.

**Figure S2.**
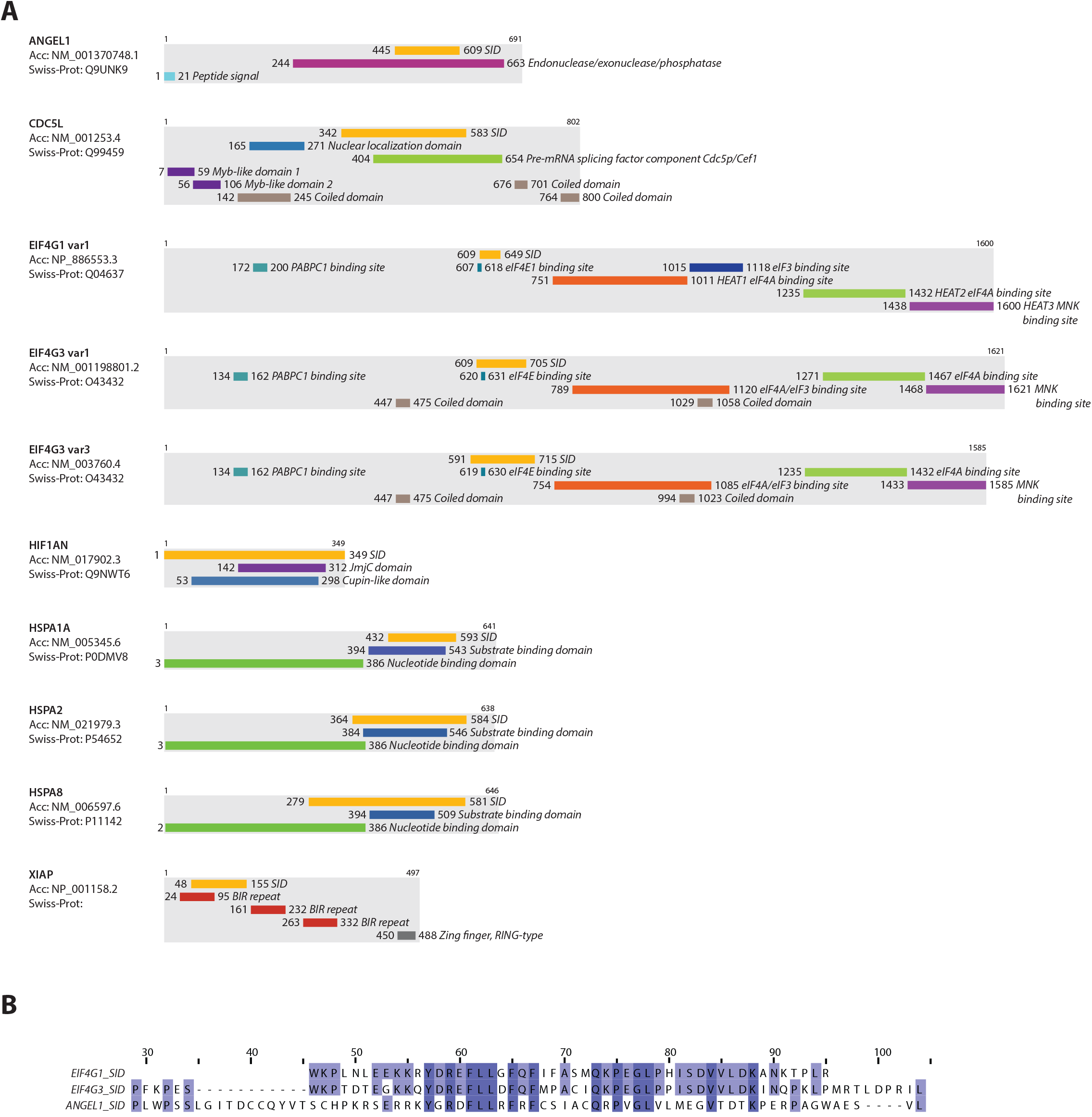
**A**. The eIF4E3 partners are represented according to their peptide chain length. The selected interaction domains (SID) as determined by Y2H, and the known functional domains of each protein are indicated. **B**. Sequence alignement of the SID identified on Angel1, eIF4G1 and eIF4G3 for eIF4E3 as represented with Jalview.

**Figure S3.**
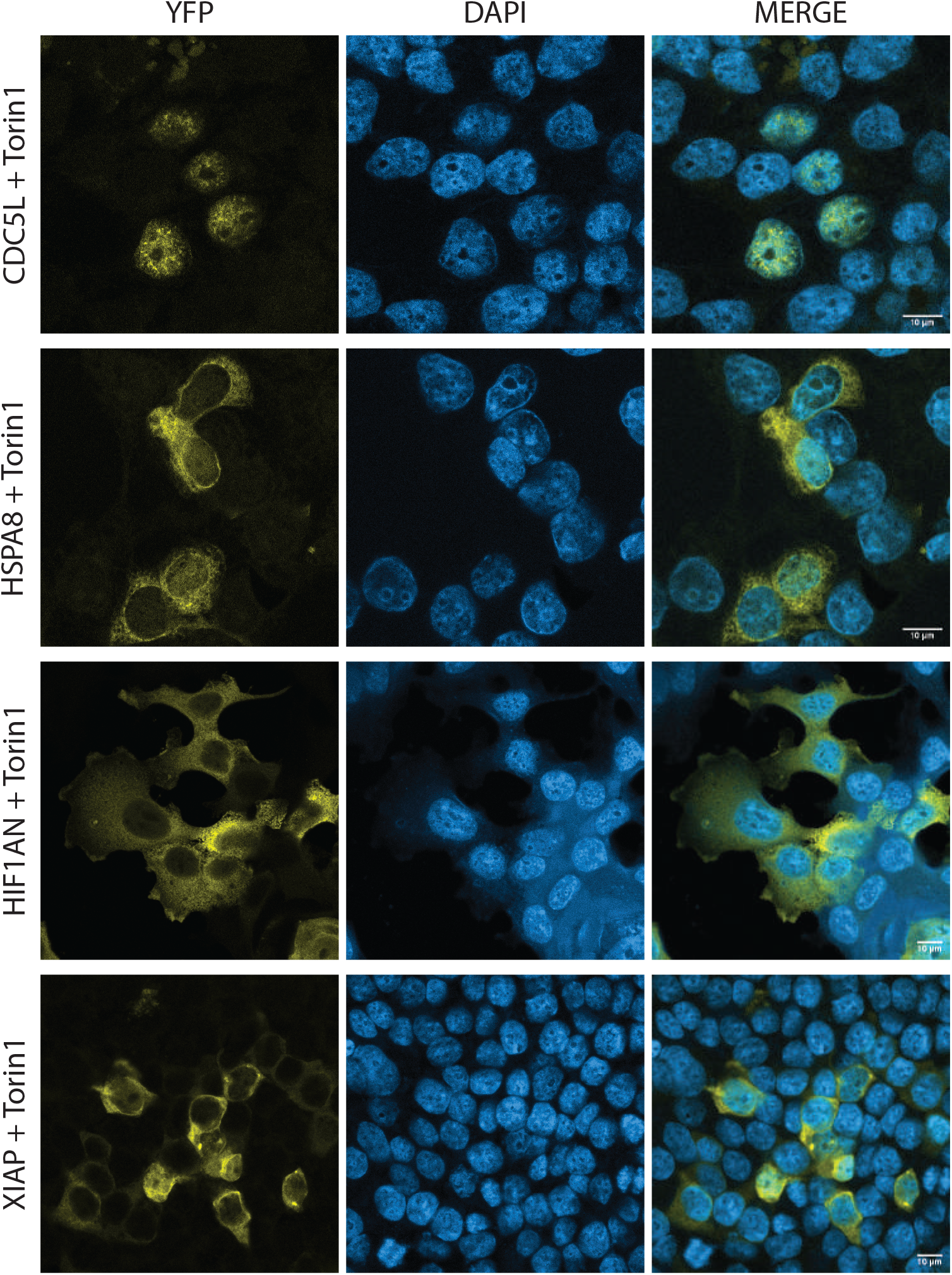
Bimolecular complementation assay using Venus YFP performed in transfected HEK293T cells treated for 2 hours with 250 nM Torin1 and monitored by confocal microscopy. Venus fragment 1 and Venus fragment 2 were respectively fused to eIF4E3 and to one of the test partners. These are indicated at the side of the left hand panel. The YTP, DAPI and merged images are shown.

**Figure S4.**
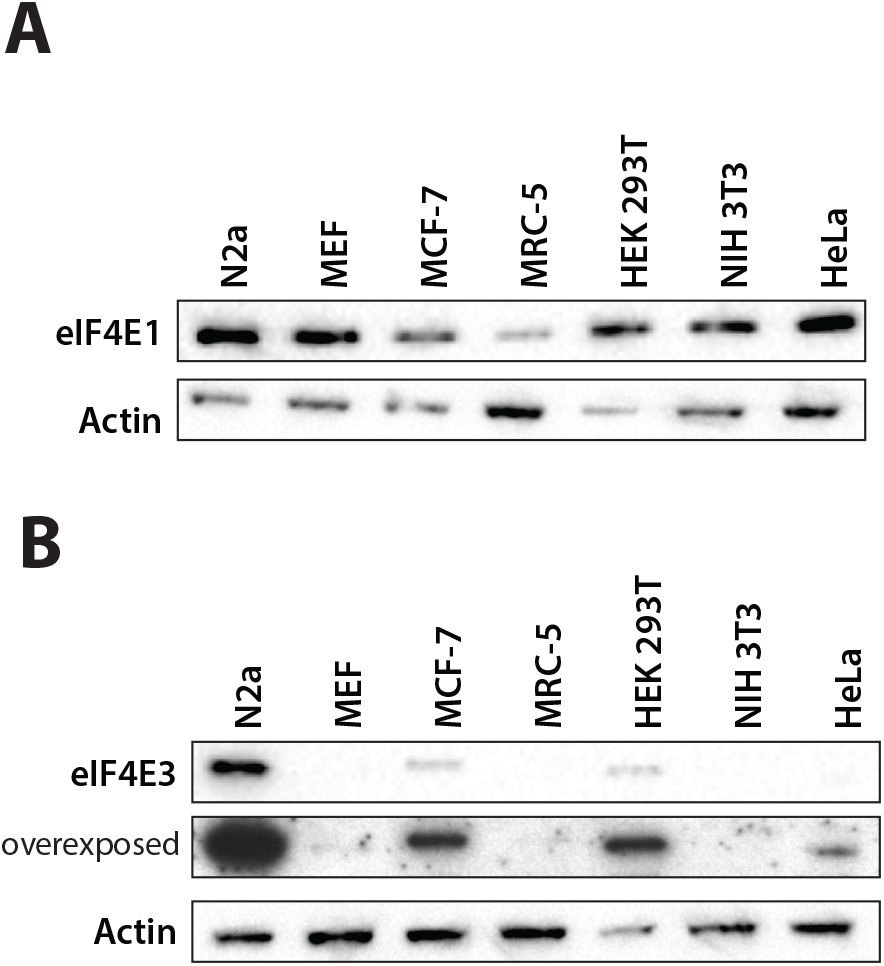
Western blots showing the level of expression of eIF4E1 (**A**) and eIF4E3 (**B**) in several mammalian cell lines.

**Figure S5.**
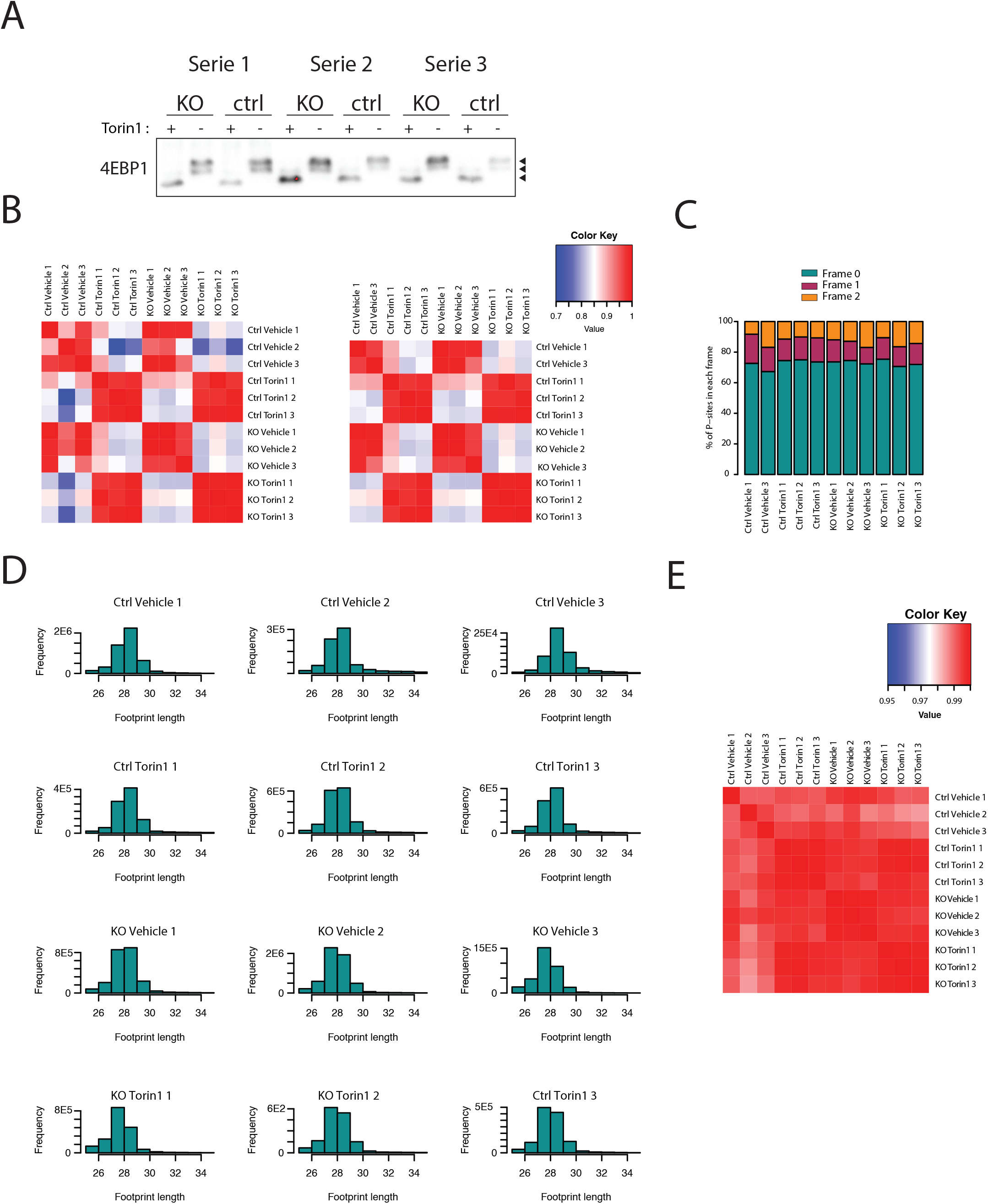
**A**. Western blot showing the phosphorylation levesl of 4EBP1 after treatment with 250 nM Torin1 in the three replicates used for ribosome profiling. **B**. Heatmap of pairwise Pearson correlations of log2 RPKMs of ribosome footprinting data. These compare all individual samples, including (**left**) and excluding (**right**) Ctrl vehicle 2. This sample was removed in all downstream analysis. **C**. Barplot of the percentage of predicted P-sites found in each frame, summed over all CDS. **D**. Histograms of ribosome footprint lengths for each sample. **E**. Heatmap of pairwise Pearson correlations of log2 RPKMs of RNASeq data for all samples.

**Figure S6.**
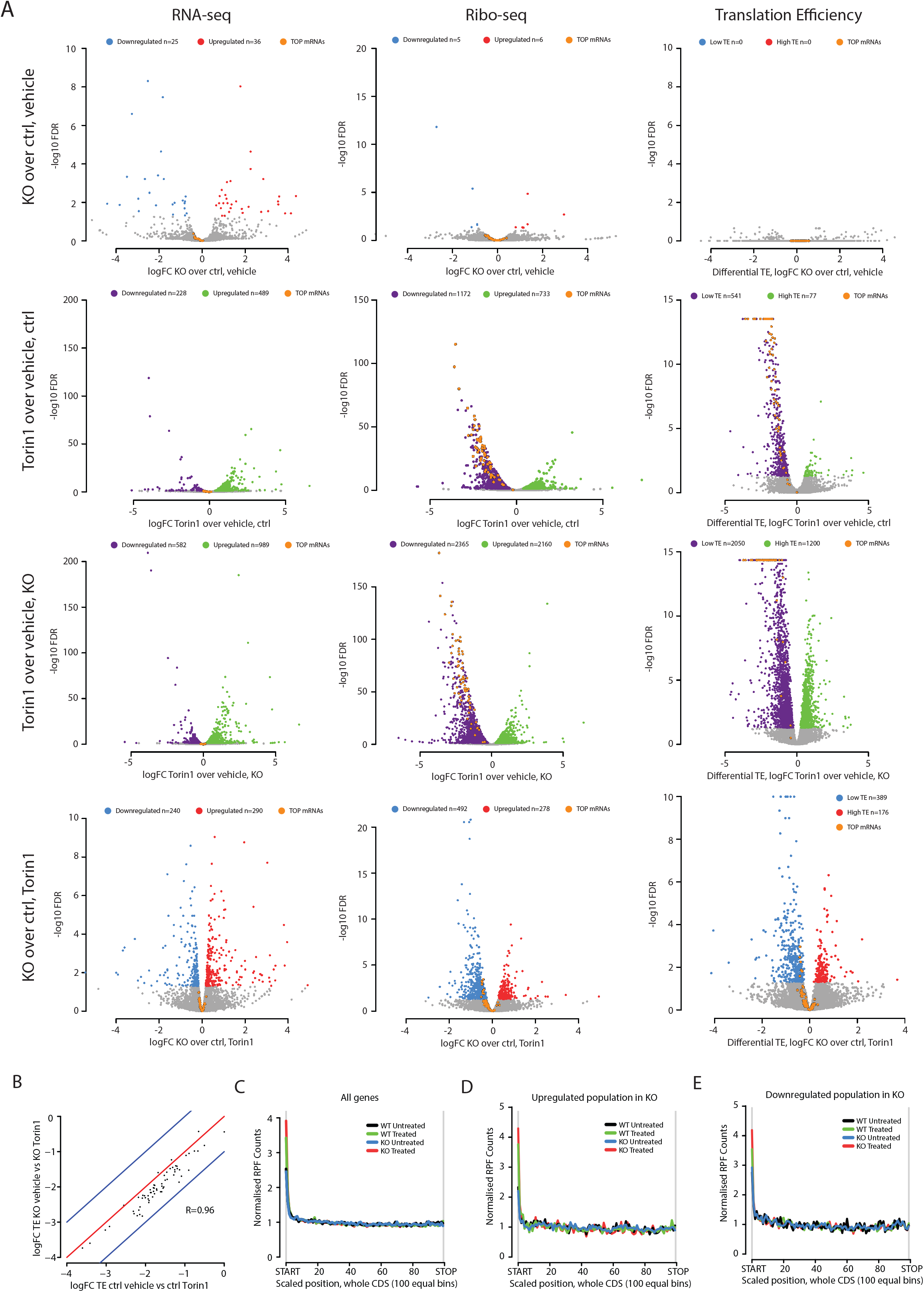
**A**. Volcano plots for all relevant pairwise comparisons of conditions (by row). The first, second and third columns show the changes for each comparison in RNA-seq, Ribo-seq and Translational Efficiency, respectively. Transcripts showing significant changes are highlighted, as are TOP mRNAs. **B.** Scatterplot of differential translational efficiency comparing vehicle and Torin1 one in ctrl vs KO. **C-E**. Scaled metagene plots for KO and WT (treated and untreated) for Ribo-seq p-site depth across the CDS for all genes (**C**) and genes upregulated (**D**) and downregulated (**E**) for RPFs in KO vs ctrl. Every CDS in the gene group was split into 100 equal bins and the number of p-sites falling in each was aggregated. These were normalised by counts for each CDS and total counts genome-wide.

**Figure S7.**
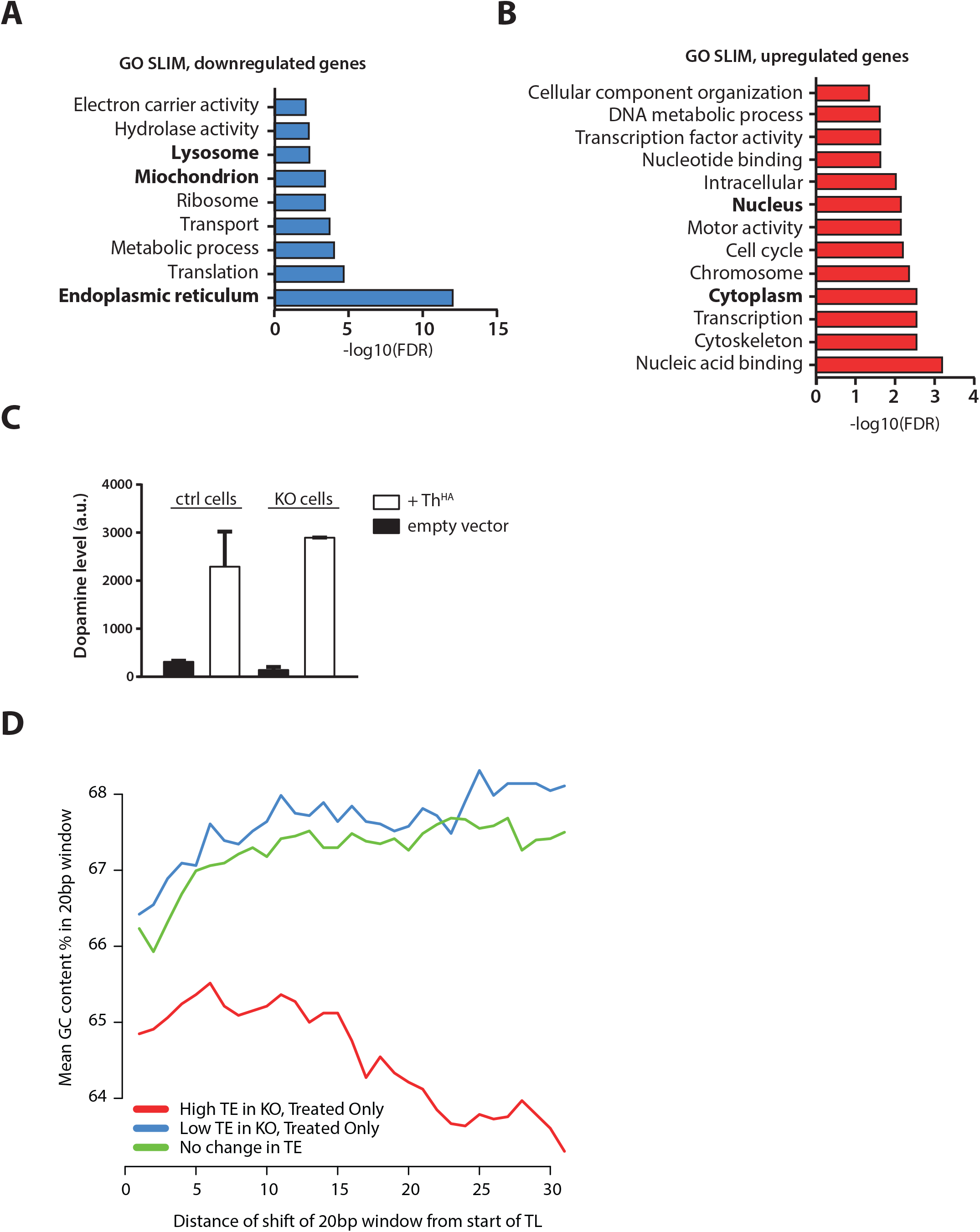
**A-B.** Barplot of FDR values for hypergeometric tests showing enrichment of various GO SLIM terms among genes downregulated (**A**) and upregulated (**B**) in KO vs ctrl, in Torin1. **C.** Histogram of the intracellular dopamine level in ctrl and KO N2a cells transduced with empty vector or with Th^HA^. **D**. Plot of mean GC percentage for 20bp windows beginning at the start of the TL and sliding base-by-base up to 30 bases from TL start. This is shown for the High TE, Low TE and No TE change groups defined in the legend of **Fig. 4E**.

